# Schlafen 12 restricts HIV-1 latency reversal by a codon-usage dependent post-transcriptional block in CD4+ T cells

**DOI:** 10.1101/2022.11.09.512793

**Authors:** Mie Kobayashi-Ishihara, Katarína Frazão Smutná, Florencia E. Alonso, Jordi Argilaguet, Anna Esteve-Codina, Kerstin Geiger, Meritxell Genescà, Judith Grau-Expósito, Clara Duran-Castells, Selina Rogenmoser, René Böttcher, Jennifer Jungfleisch, Baldomero Oliva, Javier P. Martinez, Manqing Li, Michael David, Makoto Yamagishi, Marta Ruiz-Riol, Christian Brander, Yasuko Tsunetsugu-Yokota, Maria J. Buzon, Juana Díez, Andreas Meyerhans

## Abstract

Latency is a major barrier towards virus elimination in HIV-1-infected individuals. Yet, the mechanisms that contribute to the maintenance of HIV-1 latency are incompletely understood. Here we describe the Schlafen 12 protein (SLFN12) as an HIV-1 restriction factor that establishes a post-transcriptional block in HIV-1-infected cells and thereby inhibits HIV-1 replication and virus reactivation from latently infected cells. The inhibitory activity is dependent on the HIV-1 codon usage and on the SLFN12 RNase active sites. Within HIV-1- infected individuals, *SLFN12* expression in PBMCs correlated with HIV-1 plasma viral loads and proviral loads suggesting a link with the general activation of the immune system. Using an RNA FISH-Flow HIV-1 reactivation assay, we demonstrate that SLFN12 expression is enriched in infected cells positive for HIV-1 transcripts but negative for HIV-1 proteins. Thus, codon-usage dependent translation inhibition of HIV-1 proteins participates in HIV-1 latency and can restrict the amount of virus release after latency reversal.

## Introduction

The main obstacle in curing an established HIV-1 infection is the long-lived reservoir of latently infected CD4+ T cells^1^. These cells are treatment-resistant and therefore enable the persistence of HIV-1 proviruses despite combination antiretroviral therapy (cART) and antiviral immune responses. Once cART is interrupted, HIV-1 rapidly rebounds from the viral reservoir even after years-long treatment periods^1^. During treatment, the reservoir is maintained by T cell expansion that can be activated by (i) antigen-driven proliferation, (ii) integration-site-driven proliferation, and (iii) homeostatic proliferation (Reviewed in^2^,^3^). Consequently, targeting the mechanisms governing the expansion of infected T cells represents a potential treatment strategy in HIV-1 cure attempts.

Antigen-driven proliferation triggered by T cell receptor (TCR) signaling is a strong physiological inducer of CD4+ T cell expansion. While this also reactivates latent HIV-1 and thus, can be the source of viral rebound^4^,^5^, sequential waves of polyclonal T cell stimulation in the presence of cART may result in viral reservoir reduction according to the “rinse and replace” strategy^6^. Latently infected T cells may also expand clonally when virus integration takes place within cancer-associated genes that affect cell proliferation^7-10^. Given the observed enrichment for such integration events over non-cancer-associated genes, this expansion mechanism seems common^10^. Nonetheless, latency reversal depends on position effects i.e., the exact location of the provirus within the chromosome^11^ and thus, the mere expansion of infected T cell clones does not necessarily guarantee provirus transcriptional activity. Finally, infected CD4+ T cells may expand through homeostatic proliferation (HSP) driven by interleukins IL-7 and IL-15. Unlike antigen-driven proliferation, HSP allows the expansion of HIV-1-infected CD4+ T cells without activating HIV-1 expression^12^,^13^. While these conditions strongly activate STAT5 signaling^14^,^15^, the mechanisms for HIV-1 containment are unknown. Our previous work using HIV-1-infected primary CD4+ T cells maintained under HSP culture conditions suggested a post-transcriptional block as a cause of the containment^16^. Here we demonstrate that SLFN12, a member of a conserved family of proteins with antiviral activities, participates in the maintenance of HIV-1 latency and may enable to replenish the HIV-1 reservoir pool during homeostatic proliferation *in vivo*.

## Results

### *SLFN12* is differentially expressed in homeostatic proliferating primary CD4+ T cells and is a candidate for post-transcriptional blockage of HIV-1

To decipher the mechanisms that contribute to the HIV-1 refractory state in homeostatic proliferating CD4+ T cells, we analyzed differentially expressed genes in primary CD4+ T cells that were cultured either under HSP conditions or after TCR-stimulation. For this, naive CD4+ T cells were purified from peripheral blood mononuclear cells (PBMCs) of three HIV-1 uninfected blood donors (purity > 90%) and cultured in four different ways schematically shown in Fig. 1a, A-D; cultures with IL-7 and IL-15 (HSP-cultured CD4+ T cells) after stimulation via the TCR at day 12 (B; HSP+TCR) or not stimulated (A; HSP), and cultures with IL-2 after anti-CD3/CD28 activation (TCR-cultured CD4+ T cells) with a second TCR activation at day 12 (D; TCR+TCR) or without (C; TCR). Around half of the cells maintained their naive phenotype under HSP culture conditions whereas the majority of cells became memory cells under TCR conditions (Fig 1b). On day 13, RNA was isolated from the cultured cells and transcriptomes analyzed by RNA-seq (Table S1). Genes were ranked by their standard deviation across the four culture conditions. The top 2,000 variably expressed genes were classified into four main groups, referred to as clusters I to IV by k-means clustering^17^ (Fig. 1c and Table S2). Genes of cluster I (n=384) are upregulated in both HSP and HSP+TCR conditions compared to either TCR or TCR+TCR conditions. Since our previous work demonstrated that HIV-1 proviruses in HSP condition, but not TCR conditions, were refractory to TCR activation (HSP+TCR condition)^16^, cluster I is expected to contain candidate factors that contribute to HIV-1 restriction. GO analysis of the cluster I genes showed significant enrichment of the terms *immune response, cell activation* and *immune system process* (Fig. 1d). Ten of these genes were reported to be linked to HIV-1 inhibition according to the NIH HIV interactome database^18-20^ (Table S3). For example, *CD63* has been shown to repress HIV-1 infection^21-24^ and *HAVCR2* inhibits viral budding^25^ (Fig. 1e).

**Figure 1.**
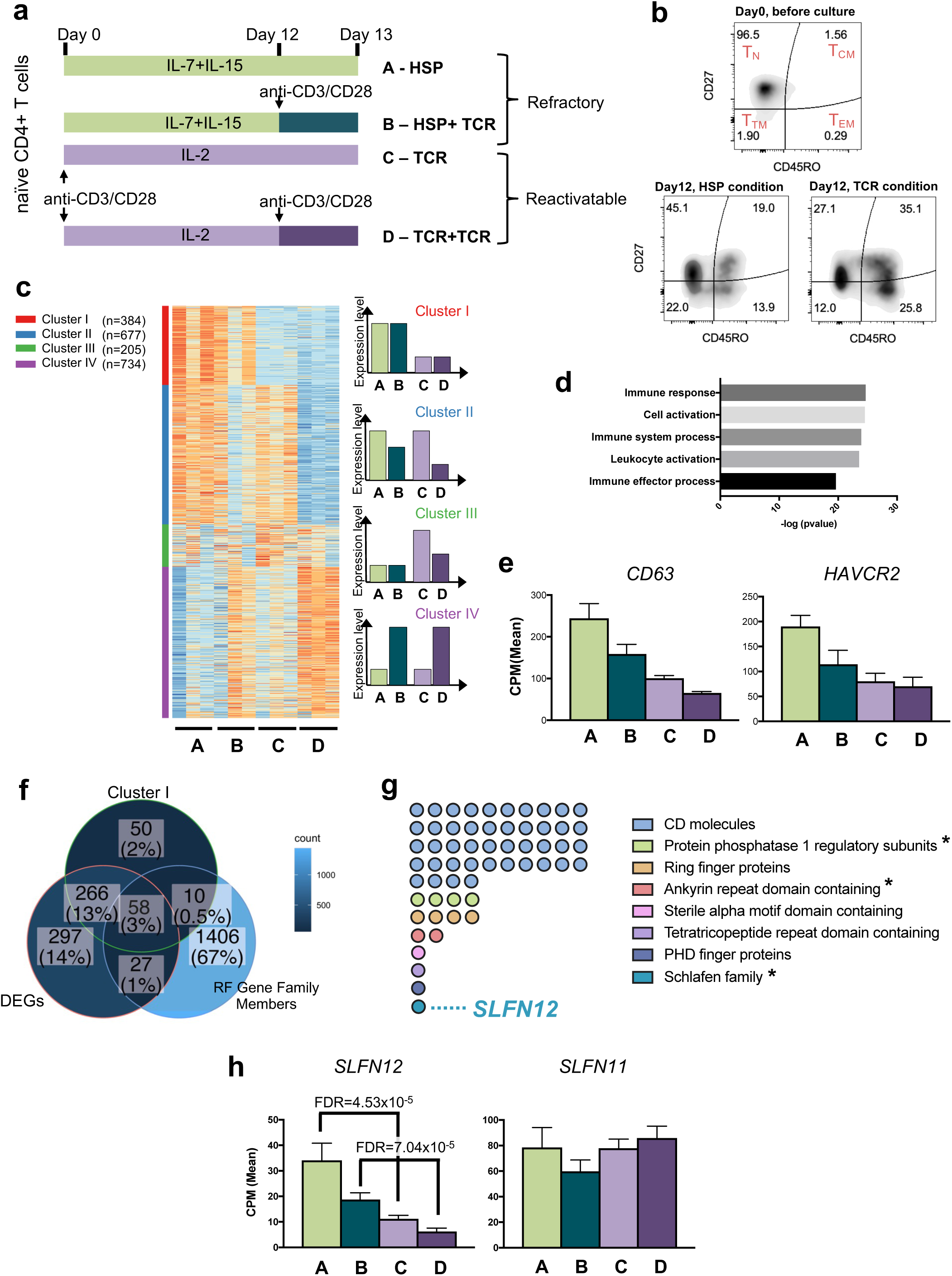
Differential RNA-seq analysis of cultured naive CD4+ T cells to identify candidate HIV-1 restriction factors. **a.** Scheme of naïve CD4+ T cell cultures. Naïve CD4+ T cells from 3 different HIV-1-negative blood donors were maintained for 13 days under 4 different conditions: IL-7+IL-15 alone (A-HSP) or with anti-CD3/CD28 activation at day 12 (B-HSP+TCR); IL-2 after anti-CD3/CD28 activation alone (C-TCR) or with a second activation by anti-CD3/CD28 at day 12 (D- TCR+TCR). On day 13, total RNA was isolated and processed for transcriptome analysis to identify candidate HIV-1 restriction factors. **b.** Phenotypes of CD4+ T cells maintained under HSP or TCR condition. A representative result of one of the three blood donors utilized in **a** is shown. Isolated naive CD4+ T cell purity was above 91%, and 96.5% for the donor shown in the upper panel. After 12-day cultivation, a naïve T cell phenotype is partially maintained in HSP conditions (CD27+ / CD45RO-, 45.1%; on average >30%), while TCR conditions increased cells with central memory phenotype (CD27+ / CD45RO+, 35.1%; on average >35%). **c.** Heatmap of the top 2,000 variable genes clustered by k-means analysis (red: upregulated, blue: downregulated). Four main clusters were identified: Cluster I, genes upregulated in both HSP and HSP+TCR conditions compared to either TCR or TCR+TCR conditions (n=384). Cluster II, genes upregulated in HSP, HSP+TCR and TCR conditions (n=677). Cluster III, genes with a trend to be upregulated in TCR condition (n=205). Cluster IV, genes upregulated by TCR stimulation at day12 (n=734). These differences in expression are visualized on the right panel for clarity. **d.** Top 5 gene ontology terms of the cluster I genes. **e.** Expression patterns of two known inhibitors of HIV-1 replication/function in the cluster I according to the NIH HIV interaction database^18-20^. Mean counts per million reads (CPM) and the standard error of the mean (SEM; n=3) are shown. **f.** Venn diagram shows overlap among (i) cluster I genes, (ii) differentially expressed genes in HSP vs TCR and in HSP+TCR vs TCR+TCR conditions (DEGs), and (iii) members of gene families that contain a known restriction factor (RF gene family members). **g.** Candidate restriction factors identified in **f** and their corresponding gene families. The asterisks show gene families that could potentially be involved in post-transcriptional and/or translational events. **h.** Expression pattern of *SLFN11* and *SLFN12* from 3 blood donors. Plots represent mean CPM ± SEM.

To further narrow down candidate restriction factors, we selected from the cluster I genes those that are (i) differentially expressed either in HSP vs TCR, or in HSP+TCR vs TCR+TCR (FDR<5%, Table S4), and that are (ii) members of a gene family that contains a known restriction factor (RF). The latter criterion was used because RF-containing gene families often have members with partially redundant functions to safeguard an organism against pathogenic threats^26^,^27^. A family member list was therefore created that includes known restriction factors^28^. It is given in Table S4. From the total of 1,501 family members, 58 genes fulfill the above criteria as candidate RFs for HIV-1 latency control (Fig. 1f). Among these 58 candidates, 44 gene products were attributed to clusters of differentiation (CD) molecules like the restriction factor CD317 (*BST2 =* Tetherin) (Fig. 1g and Table S5). However, since our previous work pointed to a post-transcriptional restriction of HIV-1 in HSP condition cultured CD4+ T cells^16^, we paid particular attention to members of three gene families that could potentially be involved in post-transcriptional and/or translational events (Fig. 1g). Protein phosphatase 1 regulatory subunits (PPP1Rs) are a gene family whose members interact with and regulate serine/threonine phosphatases. One of these is *EIF2AK2* (also known as Protein kinase R) that senses viral RNAs and enhances the integrated stress response (ISR)^29-31^, which enables the shutdown of global protein synthesis^32^. Four other family members are *PPP1R9A*, *BCL2L2*, *AATK*, and *SPRED*1 (Fig. S1). *PPP1R9A* is predicted to interact with F-actin and inhibits protein phosphatase 1-ɑ activity^33^. *BCL2L2* encodes a BCL2-like protein which is usually involved in apoptosis and thus could contribute to cell expansion under HSP conditions^34^. *AATK* is a serine/threonine-protein kinase, which is likely involved in neuronal differentiation^35^.

*SPRED1* activates MAP-kinase signaling^36^. Although these proteins have various functions, they all have the potential to deactivate translation factors by modifying their phosphorylation status in response to a viral infection. In addition, the candidate genes *UACA* and *ANKRD50* are members of the Ankyrin repeat family to which *RNASEL* (RNase L) also belongs (Fig. S1). RNase L can cleave viral RNAs to induce inflammatory responses^37^,^38^. The Ankyrin domain of RNase L is critical to forming functional dimers and sensing viral RNAs^39^. While the function of *ANKRD50* has not been well characterized, *UACA* is a known repressor of NF-*κ*B transcription^40^,^41^ and these proteins may together participate in viral RNA sensing and innate immune activation within HSP-cultured CD4+ T cells. Finally, within cluster I there was *SLFN12*, a member of the Schlafen protein family (*SLFN*). This protein has been identified in two high-throughput screenings for interferon-induced antiviral and antiretroviral factors^42^,^43^. It has moderate activity against vesicular stomatitis virus (VSV)^43^ and a mouse gammaherpes virus (MHV-68)^43^ as well as activity against several retroviruses including HIV-1^42^. Its mechanism of antiviral activity has not yet been studied. For several of the other SLFN proteins, effective antiviral functions have been reported (recently reviewed by Kim et al.^44^). SLFN11 was shown to be a restriction factor (RF) repressing HIV-1 protein translation^45^. Other human SLFN proteins namely SLFN5, SLFN13 and SLFN14 are known to attenuate the production of several viruses including influenza virus, retroviruses, and flaviviruses^46-49^. Based on these descriptions of *SLFN* genes and the statistical signal observed in our *in vitro* stimulation, we hypothesized that *SLFN12* might be another member of the *SLFN* family with an anti-HIV-1 activity that might act at a post-transcriptional level.

To first confirm *SLFN12* gene expression in CD4+ T cells under HSP and TCR culture conditions, CD4+ T cells from five additional healthy blood donors were cultured under the respective conditions (Fig. 1a) and *SLFN12* mRNAs were quantified by qPCR (Fig. S2). The expression pattern was consistent with that of the RNA-seq analysis (Fig. 1h) and differed from the other *SLFN* family mRNAs that we analyzed in comparison (Figs. S2 and S3).

### *SLFN12* inhibits HIV-1 reactivation in a post-transcriptional process

To establish a suitable *in vitro* model of viral reactivation and SLFN12 modulation, we evaluated different cell lines for *SLFN* gene expression profiles (T cell lines Jurkat, A3.01, ACH2 and embryonic kidney cells HEK 293T; Fig. 2a). *SLFN5* was expressed in all cell lines and was the only *SLFN* family member expressed in HEK 293T cells. *SLFN14* expression was not found in any of the tested cells. *SLFN11* and *SLFN12* were expressed in Jurkat, A3.01 cells and in HIV- 1 latently infected ACH2 cells. Importantly, ACH2 cells had a similar expression pattern of *SLFN11* and *SLFN12* as primary CD4+ T cells under HSP culture conditions. Therefore, given that HSP has been suggested to maintain the HIV-1 latent reservoir and that the ACH2 cell line is an established model for HIV-1 latency, we used ACH2 cells to study the effect of SLFN12 on HIV-1 reactivation. *SLFN11* was used as a positive control for the subsequent experiments.

**Figure 2.**
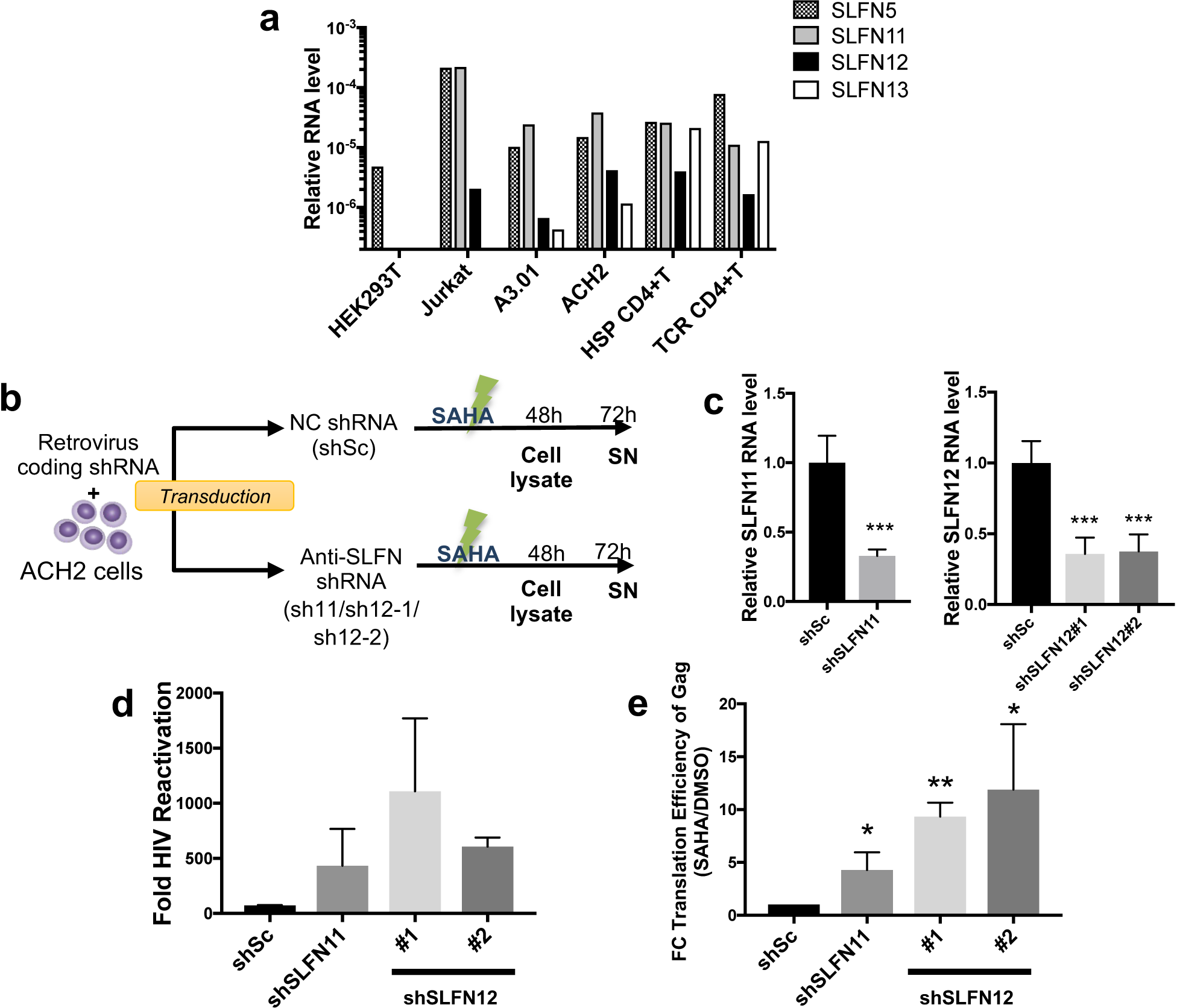
*SLFN12* affects HIV-1 latency reversal from ACH2 cells. **a.** Relative transcript levels of the human SLFN family members 5, 11, 12 and 13 in different cell lines as well as in naive CD4+ T cells under HSP or TCR culture conditions. Naïve CD4+ T cells were isolated from three independent HIV-1-negative blood donors and cultivated for 13 days with IL-7 + IL-15 (HSP condition) or with anti-CD3/CD28 antibodies + IL-2 (TCR condition) as in Fig. 1a. Transcripts were quantified by RT-qPCR with specific primer pairs and normalized to cellular *18S rRNA*. **b.** Flow chart of *SLFN*s knockdown and HIV-1 reactivation. HIV-1 latently infected ACH2 cells were transduced with retroviral vectors expressing specific shRNAs against *SLFN11* (shSLFN11) or *SLFN12* (shSLFN12#1/shSLFN12#2), or with a scrambled shRNA (shSc) as a negative control. The cells were then treated with SAHA to reactivate HIV-1. DMSO at a concentration that equals the activation mix served as the vehicle control. Cell lysates and supernatants (SN) were harvested at 48-and 72-h post-reactivation, respectively, and analyzed. **c.** Relative RNA levels of *SLFN11* (left panel) and *SLFN12* (right panel) in the knockdown ACH2 cells (n=4, mean ± SD). *** represents p<0.01 by Student’s t-test. **d.** SAHA-induced HIV-1 reactivation from ACH2 cells after *SLFN* knockdown. Fold HIV-1 reactivation was determined by titration of HIV-1-containing supernatants on TZM-bl cells and normalized to the basal HIV-1 level in DMSO-treated samples. Given is the mean ± SD for n=3. **e.** Knockdown of *SLFN11* and *SLFN12* in ACH2 cells increases translation efficiency of HIV- 1 Gag-Pr55. Plots represent fold change (FC) translation efficiency of Gag-Pr55 in SAHA-treated samples to DMSO-treated samples. Pr55 translation efficiency was calculated as a ratio of cellular Pr55 protein levels to *gag* RNA levels. The cellular Pr55 protein and HIV-1 *gag* RNA levels were quantified by Western blot and RT-qPCR, respectively. The mean FC efficiency of the control knockdown cells (shSc) was set to 1. Results show the mean ± SD of three independent experiments. Ratio paired t-test was used to calculate statical significance (*p<0.05, **p<0,005).

To knockdown *SLFN12* expression in ACH2 cells, we first generated retroviral vectors expressing specific shRNAs against *SLFN12* (shSLFN12#1/ shSLFN12#2), *SLFN11* (shSLFN11) or expressing a scrambled shRNA (shSc) as control. ACH2 cells were then transduced with these vectors and HIV-1 was reactivated by treatment with the HDAC inhibitor SAHA (also known as Vorinostat) as shown schematically in Fig. 2b. Cell lysates and supernatants (SN) were harvested at 48 and 72 hours (h) post-reactivation, respectively, and analyzed. The mRNAs of *SLFN12* or *SLFN11* were roughly 60% suppressed by the specific shRNAs (Fig. 2c). After treatment with SAHA or the vehicle control DMSO, virus-containing supernatants from ACH2 cells were titrated using TZM-bl cells. Knockdown of *SLFN12* expression resulted in at least 5.7-fold increase in infectious HIV-1 production (Fig. 2d). Moreover, the translation efficiency of HIV-1 Gag-Pr55 significantly increased (Fig. 2e, p<0.05), indicating that *SLFN12* controls HIV- 1 reactivation from latently infected T cells by repressing the translation efficiency of Gag-Pr55.

### SLFN12 restricts HIV-1 production by selectively inhibiting virus protein translation

HEK 293T cells do not express *SLFN11* or *SLFN12* (Fig. 2a). To test how SLFN12 influences HIV- 1 replication, we co-transfected HEK 293T cells with pmCherry-SLFN11 (positive control), pmCherry-SLFN12 or empty pmCherry vector (mock) together with the HIV-1 pNL-E vector, an HIV-1 provirus clone encoding *EGFP* between the *env* and *nef* coding region^50^ (Fig. 3a). Expression system of SLFN proteins was confirmed by Western blots (Fig. 3b). 48 h post-transfection, supernatants and cell lysates were collected for further analyses. SLFN12 affected HIV-1 production and strongly diminished HIV-1 titres as well as Gag-p24 protein in supernatants in a dose-dependent manner (Fig. 3c and d). This decrease was unlikely due to inhibitory effects on viral RNA processing or export (Figs. 3e, 3f, Fig. S4). Western blot analysis with specific antibodies showed downregulation of Gag-p24 and the Nef accessory protein (Fig. 3g). However, we did not observe any change in cellular GAPDH or enhanced green fluorescent protein (EGFP) derived from the pNL-E vector. Thus, SLFN12 inhibits HIV-1 production by selectively reducing translation of at least some viral proteins.

**Figure 3.**
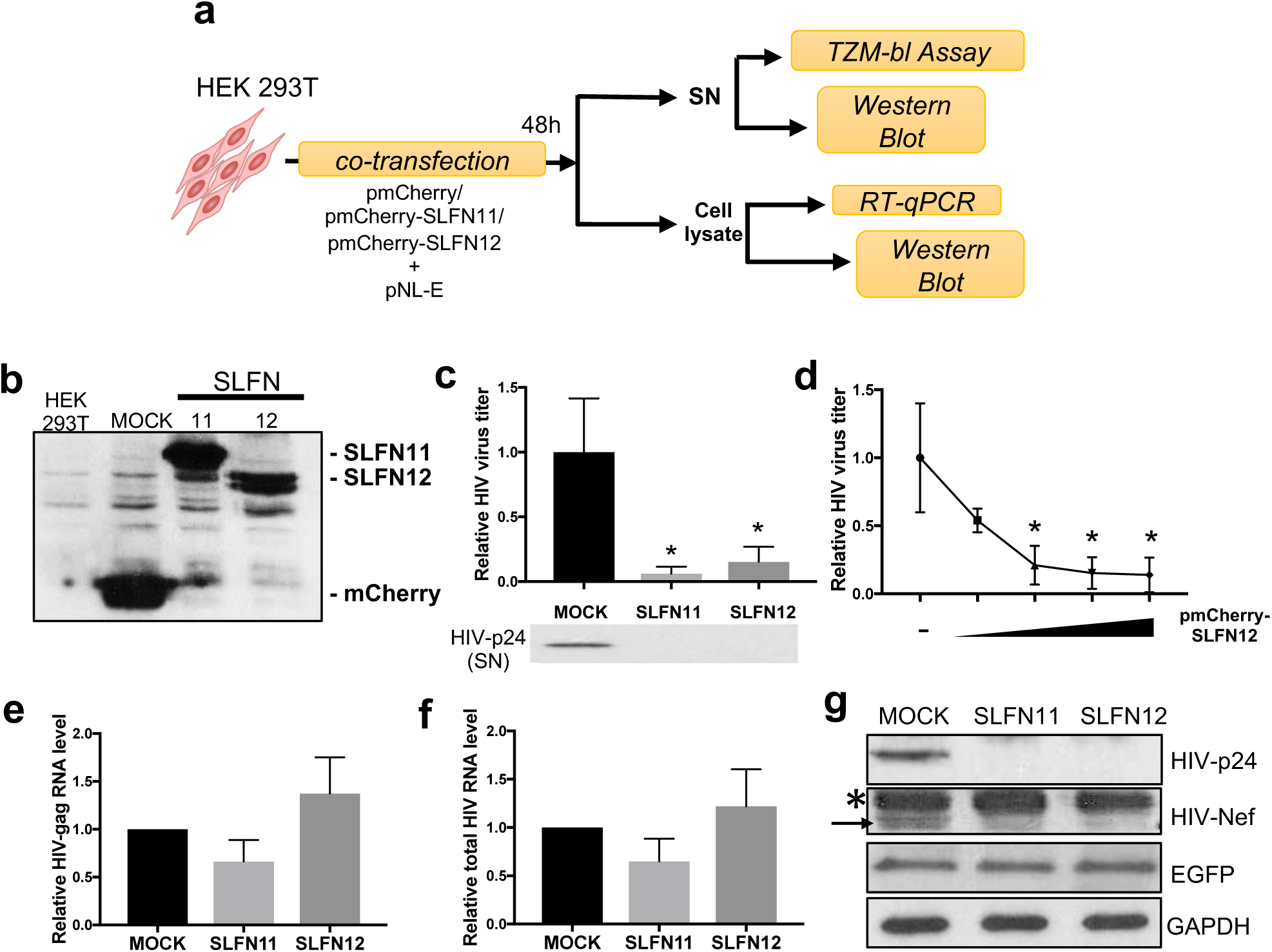
SLFN12 inhibits HIV-1 at the level of translation. **a.** Experimental outline. HEK 293T cells were co-transfected with an expression plasmid encoding mCherry-fused SLFN11 or SLFN12, or mCherry alone, and the HIV-1 vector pNL- E. At 48 h post-transfection, supernatants (SN) and cell lysates were collected and processed for further analyses as indicated. **b.** Expression of recombinant SLFN proteins in the transfected HEK 293T cells was studied by Western blots with an anti-mCherry antibody. **c.** SLFN12 expression strongly diminishes HIV-1 production from transfected HEK 293T cells. Supernatants from the co-transfected cells were titrated by the TZM-bl assay (upper panel; n=3; mean ± SD; *p<0.05 by Student’s t-test) and analyzed by Western blot (lower panel; a representative example of three independent experiments is shown). **d.** Dose-dependent inhibition of HIV-1 production by SLFN12. HEK 293T cells were co-transfected with pNL-E and increasing amounts of pmCherry-SLFN12 (0, 0.2, 0.4, 0.8 or 1.6 µg). The empty mCherry plasmid was added to maintain constant DNA amounts for all transfections. SNs were harvested 48h after transfection and titrated by TZM-bl assay (n=3, mean ± SD, *p<0.05 by Student’s t-test). **e, f.** SLFN12 expression did not significantly affect the levels of HIV-1 *gag* RNA (**e**) or total HIV-1 RNA (**f**). RNA levels were quantified by RT-qPCR and normalized to *18S rRNA* levels (n=3, mean ± SD). There were no significant differences amongst the indicated samples (Student’s t-test). **g**. Western blot from cell lysates of co-transfected HEK 293T cells with indicated antibodies. An arrow and asterisk highlight the bands of Nef and a nonspecific protein, respectively. Representative results of three independent experiments are shown.

### SLFN12 affects the translational machinery to stall ribosomes on HIV-1 mRNA

To determine how SLFN12 may influence the process of translation of viral proteins, we co-transfected HEK 293T cells with *SLFN* vectors and pNL-E, and performed a polysome profiling analysis (Fig. 4a). UV absorbance profiles showed no significant differences in total cellular RNA distribution among the three transfected cells (Fig. 4b). Given similar monosome to polysome ratios (Fig. 4c), SLFN12, as well as SLFN11, seems not to affect global mRNA translation. Next, we analyzed the distribution of *GAPDH* mRNAs, whose expression levels are not affected by SLFN12 or SLFN 11 expression (Fig. 3g), and HIV-1-*gag* mRNAs within monosome and polysome fractions. *GAPDH* mRNAs were mainly distributed in light polysomes even in the presence of the SLFN proteins (Fig. 4d, fractions 8-12, 13-17, and 18-23). However, upon SLFN12 or SLFN11 expression, HIV-1-*gag* mRNAs were shifted from light polysomes towards heavy, elongating polysomes (Fig. 4e). This increase of ribosomes per HIV- 1 *gag* mRNA, together with the observed inhibition of HIV-1-p24 expression levels (Fig. 3g), is concordant with slow or stuck ribosomes i.e., slowing down translation elongation.

**Figure 4.**
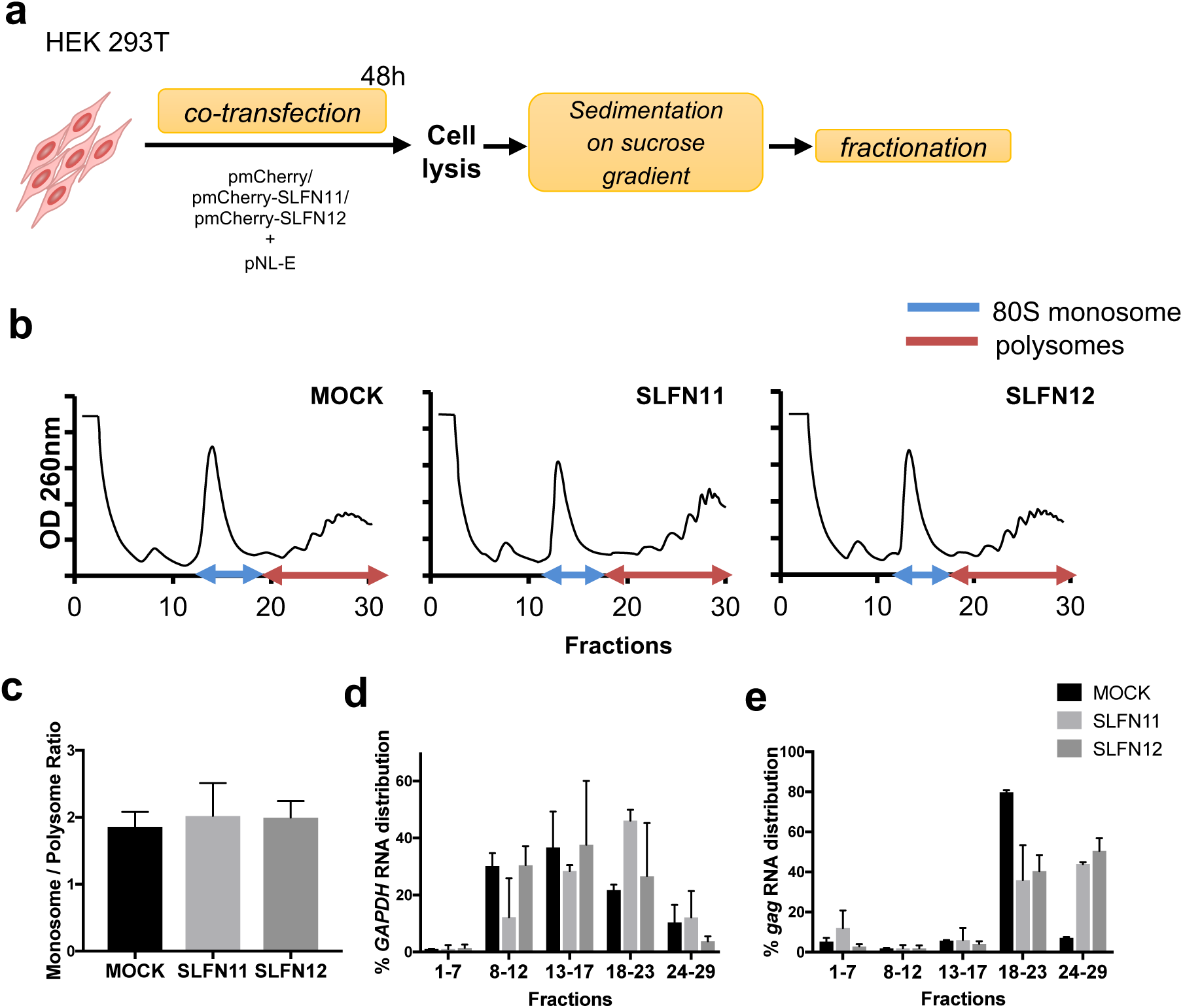
SLFN12 slows down translation elongation. **a.** Outline of polysome profiling. HEK 293T cells were co-transfected with expression plasmids encoding mCherry-fused SLFN11 or SLFN12, or mCherry alone, and the HIV-1 vector pNL-E. At 48 h post-transfection, cells were lysed and subjected to a 10-50% sucrose gradient centrifugation. The gradient was fractionated, and samples were processed for specific RT-qPCR shown in **d** and **e**. **b.** UV absorbance profiles of lysed transfected HEK 293T cells after sucrose gradient fractionation (representative result, n=3). **c.** SLFN12 expression does not influence the global ratio between monosomes and polysomes. **d, e.** SLFN12 expression shifts the HIV-1 RNA distribution towards heavy polysomes. Distribution of *GAPDH* RNA (**d**) or HIV-1 *gag* RNA (**e**) in different fractions of HEK 293T cells expressing SLFN11 or SLFN12. Pooled fractions represent free proteins (mRNPs fraction; 1-7), single ribosomal subunits (40S+60S; 8-12), 80S monosome (13-17), light-(18-23) and heavy-polysomes (24-29). The total amount of the RNAs in all fractions was set to 100%. Error bars represent standard deviations (mean ± SD; n=2).

### Translational block by SLFN12 is codon-usage-dependent

The translation elongation rate is severely affected by codon optimality of transcripts and the availability of cognate transfer RNAs (tRNAs)^51^,^52^. SLFN11 has been shown to inhibit HIV-1 translation in a codon-usage-dependent manner^45^. To test whether also SLFN12 affects HIV-1 translation elongation in a codon-usage-dependent manner, we first compared viral codon frequencies to RefSeq-based human coding sequences using the Codon Adaptation Index (CAI), a general metric to analyze codon usage bias^53^. As previously described^54^, HIV-1 sequences harbor less optimal codons (lower CAI values) compared to most human transcripts including *GAPDH* (Fig. 5a). Next, we compared the effect of SLFN12 on Gag-p24 protein production from wild-type (wt) and codon-optimized HIV-1-*gag* transcripts. For this, HEK 293T cells were co-transfected with the SLFN12 expression vector plus a vector expressing either HIV-1-*gag wt* (pGag-wt) or a codon-optimized HIV-1-*gag* (pGag-opt). Protein and mRNA levels were then analyzed 48 h later (Fig. 5b). Compared to mock, SLFN12 strongly diminished translation of Gag-p24 protein transcribed from HIV-1-*gag wt* (Fig. 5c), however, protein expression in cells transfected with pGag-opt remained unchanged (Fig. 5d). Since the RNA levels of HIV-1-*gag wt* and HIV-1-*gag opt* were not significantly altered by SLFN12, HIV-1 inhibition by SLFN12 (Fig. 3) may indeed occur at a post-transcriptional step.

**Figure 5.**
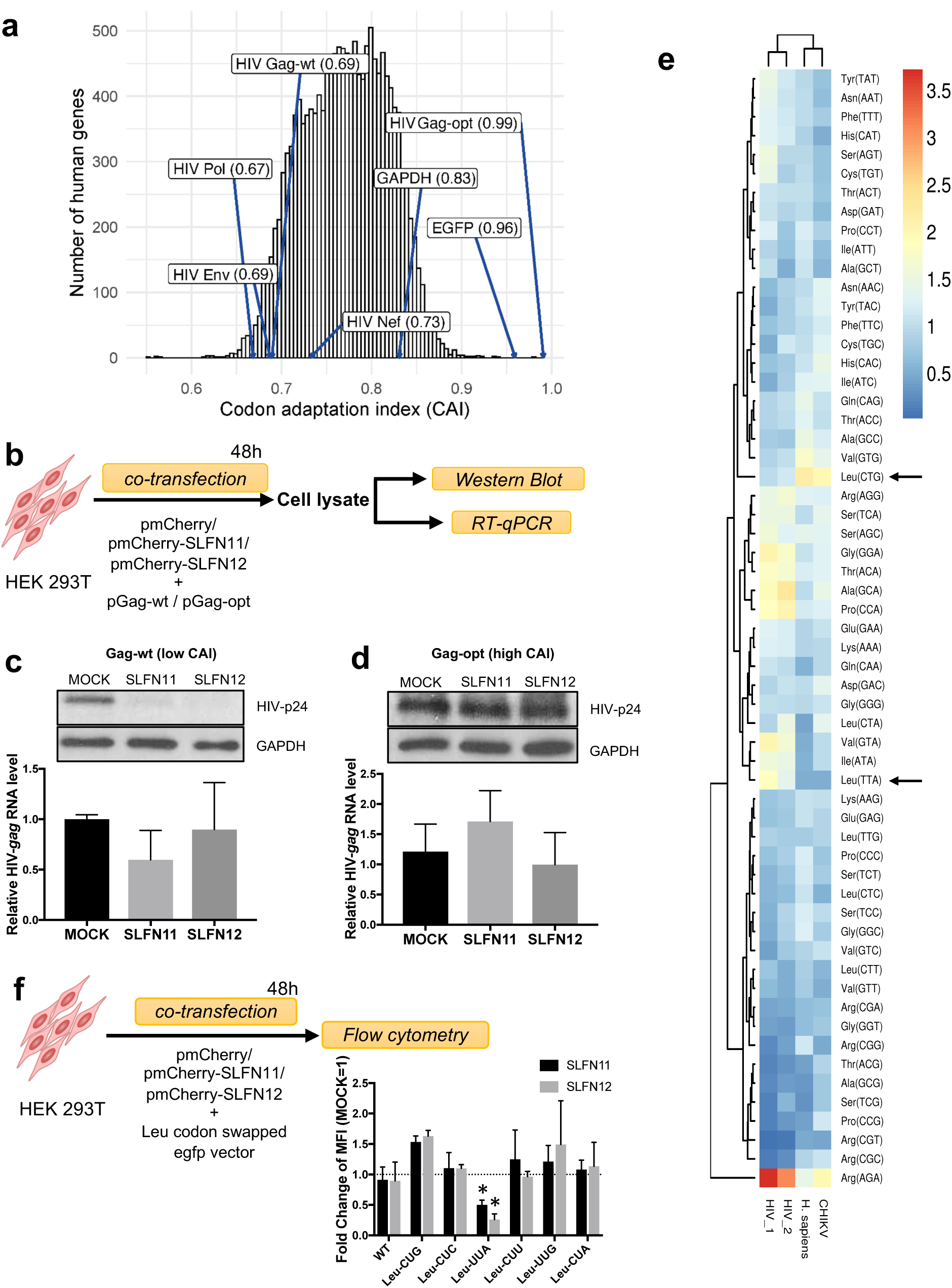
SLFN12-mediated inhibition of HIV-1 translation is codon-usage dependent. **a.** Histogram of codon adaptation indices (CAIs) for the human RefSeq transcript hg38. CAIs of HIV-1 sequences, *GAPDH* and EGFP are highlighted for comparison. The mean CAI of human transcripts was 0.77. Gag-opt, codon-optimized gag sequence as in plasmid pGag-opt. **b.** Experimental outline. HEK 293T cells were co-transfected with expression plasmids encoding mCherry-fused SLFN11 or SLFN12, or mCherry alone, and an HIV-1 vector expressing Gag-p24 with either wild-type (pGag-wt) or optimized codon usage (pGag-opt). At 48 h post-transfection, transfected cells were lysed and analyzed by Western blot and RT-qPCR. **c, d.** Upper panel: Western blot detection of p24 from HEK 293T cells expressing wild-type Gag (Gag-wt, c) or codon-optimized Gag (Gag-opt, d). Lower panel: Relative HIV-1-gag RNA levels from indicated expression vectors (n=3, mean ± SD). There were no significant differences between the RNA levels amongst the indicated samples (Student’s t-test). **e.** Heatmap of relative synonymous codon usage of human (H. sapiens), HIV-1, HIV-2 and Chikungunya virus (CHIKV) transcripts. Codons Leu-UUA (Leu(TTA) in the graph) and Leu-CUG (Leu(CTG)) are highlighted with arrows. **f.** SLFN11 and SLFN12 specifically inhibit Leu-UUA codon-swapped EGFP expression (n=3, mean ± SD). The upper panel shows the outline of the experiment. Wild-type EGFP expression vector (WT) or Leu codon-swapped vectors (Leu-CUG, Leu-CUC, Leu-UUA, Leu-CUU, Leu-UUG, and Leu-CUA) were individually transfected together with mCherry (MOCK), mCherry-fused-SLFN11 or 12 expression vector into HEK 293T cells. After 48h, relative mean fluorescence intensities (MFI) were measured by flow cytometry. The fold change MFI values are shown; the MFI of mock transfection was set to 1. The asterisks represent statistical significance (*p<0.02, by one sample t-test).

To identify which codons may de-optimize the HIV-1 codon usage, we compared the relative synonymous codon usage (RSCU)^55^ amongst HIV-1, HIV-2, Chikungunya virus (CHIKV) as an example of an unrelated virus, and humans (*H.sapiens*). As previously reported^56^,^57^, HIV-1 and -2 utilize many A-ending codons that are rare in humans and CHIKV (Fig. 5e and Table S6). Among A-ending codons, both HIV-1 and HIV-2 use Leu-UUA very frequently instead of Leu-CUG that is common in humans. To test whether Leu codon swapping within *EGFP* would render its expression sensitive to SLFN12 inhibition, HEK 293T cells were co-transfected with the SLFN12 expression vector plus synonymous *EGFP* constructs differing in their codon usage. *EGFP* expression was analyzed by flow cytometry 48 h later (Fig. 5f). SLFN12 inhibited only Leu-UUA swapped *EGFP*. Altogether our results demonstrate that the attenuation of HIV-1 protein production by SLFN12 is codon-usage dependent.

### SLFN12-mediated HIV-1 suppression depends on a putative tRNase cleavage domain

To better understand the mechanism by which SLFN12 may interfere with HIV-1 protein translation, we concentrated on the structural features of SLFN protein family members. The structure of SLFN12 was recently resolved by cryo-EM and suggested an RNase activity^58^. SLFN11 and SLFN13 are known to degrade tRNAs^49,59-61^. Comparison of sequences and conformations of SLFN11, 12 and 13 showed a 34.8% similarity between SLFN12 and SLFN11 or SLFN13, and a 75.8% similarity between SLFN11 and SLFN13. Protein backbone structures were similar between the 3 proteins except that SLFN12 had a short C-terminal domain (Fig. 6a). Based on these structural features, we hypothesized that SLFN12 may cleave the rare Leu-UUA tRNA and reduce its cellular concentration.

**Figure 6.**
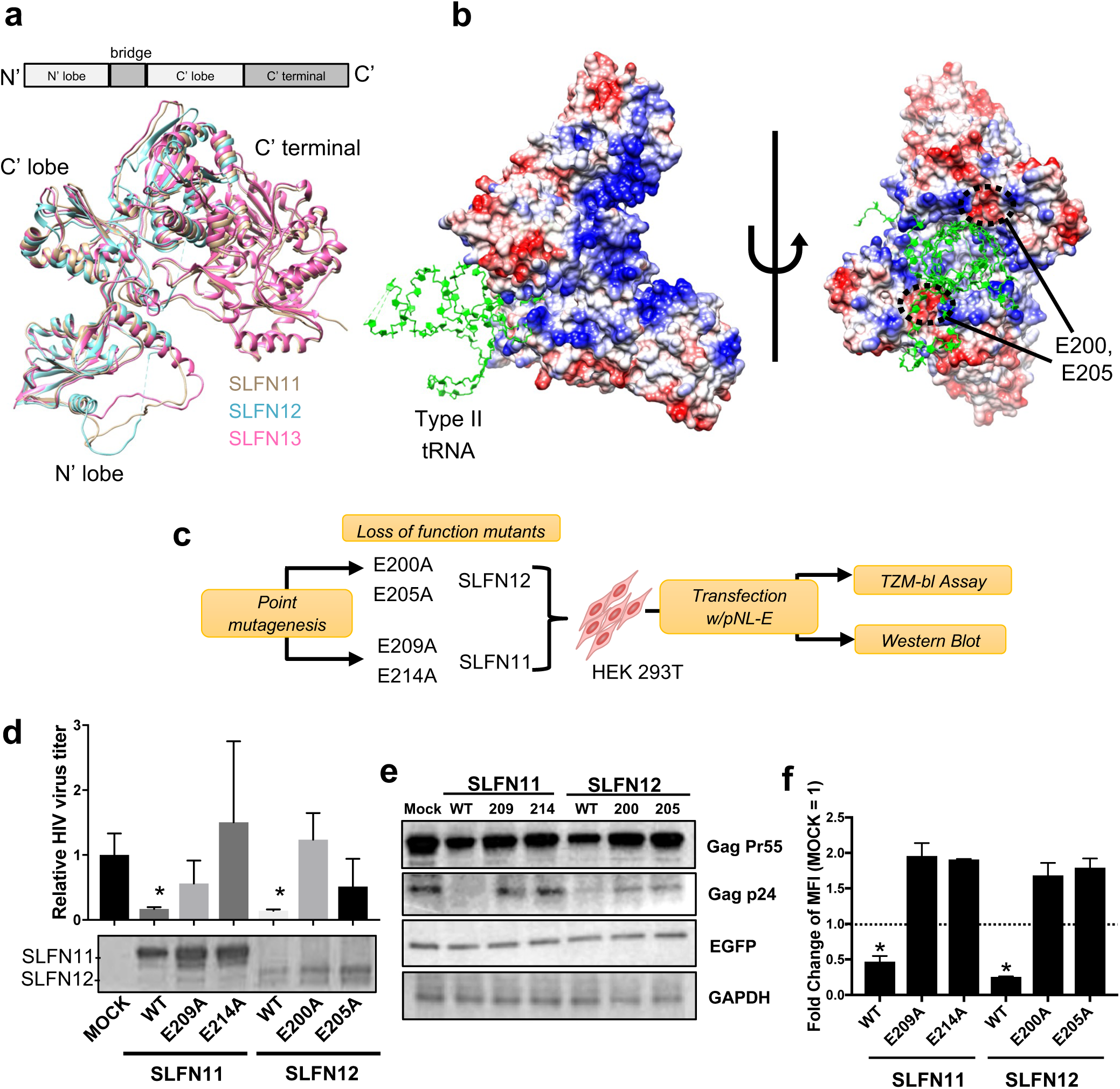
SLFN12-mediated HIV-1 suppression depends on a putative tRNase cleavage domain. **a.** Conformational similarity of human SLFN11 (beige), SLFN12 (cyan) and SLFN13 (magenta). The cryo-EM structure of SLFN12 (7LRE) and alpha-fold predicted SLFNs 11 (Q7Z7L1) and 13 (Q68D06) were downloaded from Uniplot and compared by Chimera software (version 1. 13. 1). SLFN12 has a short C-terminal domain. **b.** Structural models of SLFN12 dimer / tRNA complexes. The cryo-EM structure of SLFN12 dimer (7LRE) is shown as surface representations with positively (red) and negatively (blue) charged areas. A human type II tRNA (selenocysteine tRNA, 3HL2) is docked with the SLFN12 dimer based on the predicted rat SLFN13-tRNA_49_. Putative catalytic sites glutamic acid 200 (E200) and 205 (E205) for SLFN12. They are presented inside the black circles. **c.** Experimental outline. mCherry-fused SLFN expression vectors or their respective active site mutants E209A and E214A of SLFN11, and E200A and E205A of SLFN12 were individually co-transfected with HIV-1 pNL-E vector into HEK 293T cells. 48 h post-transfection, cells and supernatants were harvested and processed for further analyses as indicated. The following results are from three independent experiments (d, e). **d.** Mutants of the tRNase cleavage sites of SLFN12 lose their anti-HIV-1 activity (mean ± SD). Asterisks indicate significant differences compared to MOCK-transfected cells calculated by Student’s t-test. The lower panel shows expression levels of the SLFN proteins in a Western blot using an anti-mCherry antibody. **e.** Western blot showing intracellular HIV-1 Gag Pr55 and p24 levels in SLFN-expressing cells. EGFP and GAPDH were used as controls. **f.** The Leu-UUA codon swapped vector was transfected with indicated SLFN mutant expression vectors into HEK 293T cells. After 48h, relative mean fluorescence intensity (MFI) was measured by flow cytometry. The fold change MFI values are shown; the MFI of mock transfection was set to 1. Asterisks indicate decreased values with statical significance (*p<0.05 by Student’s t-test; n=2; mean ± SD).

Leucine tRNAs are categorized as type II tRNAs that have an expanded variable loop compared to type I tRNAs^62^. We, therefore, simulated docking between the SLFN12 dimer and human type II tRNA structure available in the Uniplot database (Fig 6b). A type II tRNA potentially fits into a space formed by the SLFN12 dimer. Furthermore, putative catalytic sites, which are composed of the negatively charged glutamic acids E200 and E205 in SLFN12, mapped closely to the docked tRNA. These putative RNase active sites were conserved as E209 and E214 in SLFN11^61^. Thus, SLFN12 may exert its codon-specific HIV-1 inhibitory effect through the two putative cleavage sites E200 and E205. To investigate this hypothesis, we generated expression vectors of SLFN11 and 12 in which the respective active site glutamic acids were converted to alanine. HEK 293T cells were then co-transfected with the individual *SLFN* constructs and pNL-E, and the antiviral activity was tested by TZM-bl assays and Western blot (Fig. 6c). All active-site mutants of SLFN11 and SLFN12 reduced SLFNs anti-HIV-1 activity (Fig. 6d) and Gag-p24 translation inhibition (Fig. 6e). Likewise, these mutants lost their codon-dependent inhibition of Leu-UUA swapped-*EGFP* expression as shown by the increase of mean fluorescence intensities of HEK 293T cells co-transfected with Leu-UUA swapped-*EGFP* plus the individual *SLFN* mutants (Fig. 6f). Together these results demonstrate that the putative tRNA cleavage sites E200 and E205 of SLFN12 mediate the post-transcriptional, codon-dependent blockage of HIV-1.

To test whether SLFN12 degrades tRNAs, we quantified tRNAs in SLFN-transfected cells by RNA gel electrophoresis. Visual inspection of the ethidium bromide-stained polyacrylamide gel showed a slight decrease in type II tRNA abundance (Fig. S5a). Likewise, type I and II tRNA quantification by image J normalized to 5.8S rRNA levels showed a decrease in type II tRNA by SLFN12, although with less activity than the positive control SLFN11 (Fig. S5b).

Involvement of SLFN12 in HIV-1 post-transcriptional restriction within infected individuals

To study SLFN12 expression and its putative role in the control of HIV-1 latency in HIV-1- infected individuals, we first analyzed *SLFN12* expression in PBMCs from HIV-1 high (High, n=16) and low viremic (Low, n=30) individuals, viremic controllers (VC, n=11), and elite controllers (EC, n=12). *SLFN12* mRNA expression levels per CD4 count showed significant differences among each group, and a tendency to increase with disease progression (Fig. 7a). The *SLFN12* mRNA level was also positively correlated with viral RNA and DNA level, respectively, indicating that *SLFN12* is expressed in response to viral load (Fig. 7b and c). To further test whether SLFN12 may be involved in post-transcriptional HIV-1 latency control *in vivo*, patient’s PBMCs were treated with or without LRAs (Romidepsin and Ingenol), stained with anti-Gag-p24 and anti-SLFN12 antibodies plus anti-HIV-1-RNA probes, and analyzed by FISH-Flow assays as previously described^63^. The gating strategy is shown in Fig. 7d. SLFN12 expressions in p24 positive(+), HIV-1-RNA+ / p24 negative(−), and both negative populations were characterized (Fig. 7e and f). SLFN12-expressing cells were enriched in the population of HIV-1-RNA+ / p24 – cells, which is consistent with a role in post-transcriptional HIV-1 latency control.

**Figure 7.**
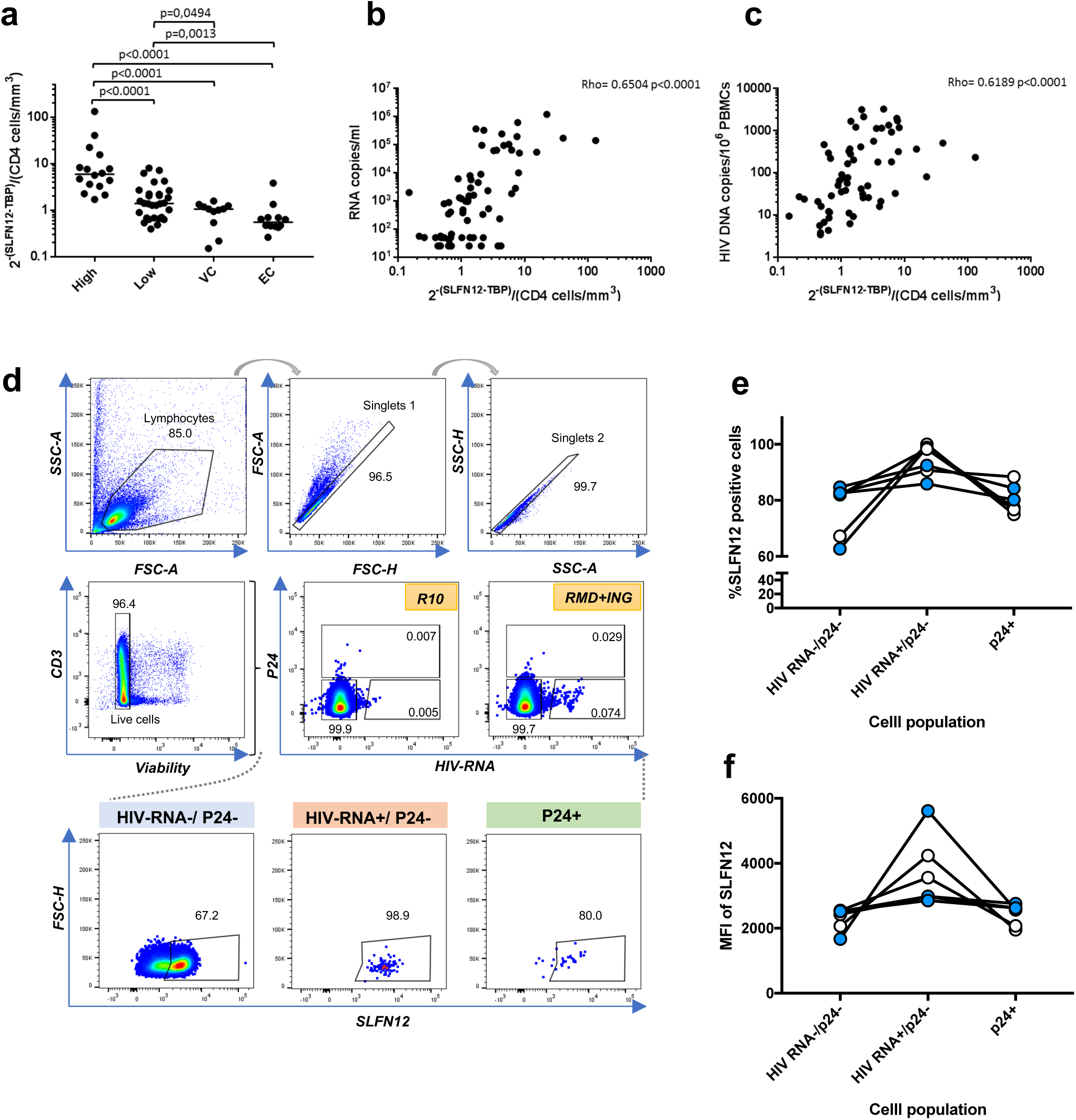
Involvement of *SLFN12* gene in HIV-1 post-transcriptional restriction in patients. **a.** Expression level of *SLFN12* in PBMCs from HIV-1-infected individuals. SLFN12 RNA levels were quantified using PBMCs from HIV-1 high viremic- (High, n=16) and low viremic- (Low, n=30) patients, and viremic- (VC, n=11) and elite-controllers (EC, n=12). SLFN12 levels were normalized by TBP RNA levels and CD4+ T cell counts per µl. Each point represents the values of a single individual. p-values were calculated by Mann-Whitney test. **b.** Correlation between *SLFN12* expression levels and viral RNA loads. Each point represents the values of a single individual. The correlation was calculated by Spearman’s rank test. **c.** Correlation between *SLFN12* expression levels and HIV-1 provirus loads. Each point represents the values of a single individual. Spearman’s rank test was applied to analyze the correlation. **d-f.** SLFN12 expressing CD4+ T cells are enriched in HIV-1 RNA+/p24- cells. Gating strategy to measure SLFN12 expression in patient CD4+ T cells (d). Gating strategy consisted of selecting lymphocytic cells by FSC- and SSC-scatter, followed by a double doublet exclusion, dead cells exclusion and finally an HIV-1-RNA+ and p24+ gate from where SLFN12+ cells were identified. The proportion of SLFN12 expressing cells (e) or relative MFI of SLFN12 (f) in p24-/HIV-1 RNA-, p24-/HIV-1 RNA+, or p24+ cells. Blue circles, patient samples treated with Romidepsin (RMD) plus Ingenol (ING) *ex vivo* (n=3); White circles, untreated patient samples (R10 in d, n=3).

## Discussion

The factors and mechanisms that maintain HIV-1 latency and thus hamper antiviral cure strategies are incompletely understood. Here we demonstrate that SLFN12, a member of the conserved family of Schlafen proteins, is a novel HIV-1 restriction factor that inhibits HIV-1 translation in a codon-usage dependent manner and participates in virus containment in homeostatically proliferating CD4+ T cells. The mechanism of action of SLFN12 was similar to that of the previously described SLFN11, despite a limited sequence homology between the two Schlafen family members. As the two genes showed differences in their transcription patterns, their antiviral functional redundancy may indicate a way to ensure virus control under different cell growth conditions.

Homeostatic proliferation is a physiological process that restores the peripheral T cell pool after lymphopenia. It requires the cytokines IL-7 and IL-15, and HLA-restricted T cell receptor triggering of low affinity^64^. Importantly, these conditions allow HIV-1-infected T cells to expand without virus production^12^,^13^. Our previous observations with latently infected primary CD4+ T cells under HSP culture conditions^16^, and that of Mohammadi et al. using another primary CD4+ T cell model of HIV-1 latency^65^, suggested that virus production from provirus-containing cells can be restricted by a post-transcriptional rather than a transcriptional block. Our work here now suggests that SLFN12 is an important component of this post-transcriptional block. It may enable HIV-1-infected cells to remain undetectable to adaptive immune responses and antiviral therapies because the underlying effector mechanisms require viral protein expression for exerting their antiviral function. Consequently, blocking SLFN11 and SLFN12 function and thus, increasing viral protein production would help expose HIV-1 infected cells more effectively to the immune system and antiviral drugs. This should lead to increased HIV- 1-infected cell elimination by the immune system and the reduction of the latent reservoir of infected cells *in vivo*. Studies in this direction are clearly worth pursuing.

Functionally, SLFN gene members have multiple and divergent biological roles in host defense^45-49^, cell differentiation^66^,^67^ and cancer drug sensitivity^58,59,68,69^. SLFNs 5, 11, 13 and 14 have been described as virus restriction factors showing activities against diverse RNA and DNA viruses while SLFN12 until now was considered a candidate antiviral protein based solely on the result of high-throughput screens for interferon-induced antiviral factors^42-44^. Our present work here extends these studies and demonstrates the mechanism by which SLFN12 inhibits HIV-1 and restricts latency reversal in a codon usage-dependent manner. Interestingly, SLFN11 and 12 showed comparable anti-HIV-1 properties that in both cases were abrogated by mutations in the proteinś RNase domains. However, while the C-terminal helicase domain of SLFN11 seems required for its anti-HIV-1 activity^60^, SLFN12 lacks this region and thus exerts its antiviral activity without it. Furthermore, SLFN11 also encodes a nuclear localization signal while SLFN 12 does not. Thus, the two SLFN proteins might exert their effect from both the nuclear as well as the cytoplasmic compartment of the cell. Supposing that the antiviral mechanism is mediated by the tRNA cleavage activity as demonstrated for SLFN11^59-61^ and SLFN13^49^, and suggested for 12 here as well (Figs. 6a-f), an effect in the nucleus where the tRNA is generated and in the cytoplasm where the tRNA is used for mRNA-directed translation would be most effective. However, many unknowns with respect to the regulation of the RNase activities of SLFN11 and SLFN12 as well as their possible individual and combined effects on the tRNA pool remain and require further studies.

Our data from a cohort of HIV-1-infected individuals showed a positive correlation between *SLFN12* expression and HIV-1 viral load (Figs. 7a-c), suggesting a link with the general activation of the immune system. Indeed, *SLFN* family gene expression is considered interferon-inducible^42,43,70,71^. Furthermore, our transcriptome-based identification of *SLFN12* was derived from the screening of homeostatic proliferating primary CD4+ T cells *ex vivo* (Fig. 1). This mode of proliferation requires the cytokines IL-7 and IL-15 whose expression *in vivo* is directly linked to reduced CD4+ T cell counts, and high HIV-1 loads^72-74^. Interestingly, *SLFN12* was just one amongst several candidate HIV-1 restriction factors induced under these conditions (Figs. 1c-g), thus suggesting that CD4+ T cells during homeostatic proliferation expand highly protected.

To finally evaluate whether SLFN12 restricts HIV-1 protein expression at a post-transcriptional level *ex vivo*, we combined FISH-Flow assays to detect intracellular HIV-1 RNA with anti-Gag- p24 and anti-SLFN12 antibody staining and analyzed PBMCs of several infected individuals. A trend towards the enrichment of SLFN12 expression within HIV-1-RNA+/p24– cells was observed in these patients. The same was true when considering the mean fluorescence intensities of SLFN12-expressing cells. This supports our *in vitro*-experiments with HIV-1- infected cell lines and suggests that SLFN12 also restricts the generation of infectious HIV-1 in people living with HIV (PLWH). However, the frequency of HIV-1 RNA+ and/or p24+ CD4+ T cells within the PBMCs of HIV-1-infected individuals was low (0.01∼0.1%), and the current FISH-flow analysis is technically limited in that it cannot discriminate whether infected cells containing complete and competent HIV-1 proviruses from those with defectives forms. Therefore, our observations should be interpreted with caution.

In conclusion, SLFN12 now adds to the growing number of restriction factors that a host utilizes to downregulate HIV-1 production. Importantly, by blocking virus protein production at a translational level, SLFN11 and 12 also help HIV-1 and infected cells to escape from antiviral therapy and avoid immune-mediated destruction, facilitating the virus to reside within its host organism. This makes SLFN11 and 12 interesting therapeutic targets in strategies aiming to eliminate HIV-1 persistence. An experimental examination of this interesting hypothesis clearly deserves further consideration.

*Note added in proof.* While preparing our manuscript, Lee et al published that SLFN12 selectively cleaves Leucine UUA tRNA (*Nat.Chem.Biol.* online on 27^th^ Oct 2022). This is in complete agreement with our observations described here.

## Methods

### Ethics statement

PBMCs from healthy donors and HIV-1 infected patients were obtained from the Hospital Universitari Vall d’Hebron (PR(AG)270/2015), and IrsiCaixa (CEIC: EO-12-042 and PI-18-183) in Barcelona, Spain.

### Cell lines and PBMCs

ACH2 cells were cultured in RPMI medium (Gibco) supplemented with 10% heat-inactivated fetal bovine serum (FBS, Sigma Aldrich) and 1% of penicillin/streptomycin mix (P/S, Gibco). HEK 293T cells (ATCC) and TZM-bl cells (NIH AIDS Reagent Program) were maintained in DMEM medium (Gibco) supplemented with 10% heat-inactivated FBS and 1% P/S. All cell lines were incubated at 37°C in presence of 5% CO_2_. PBMC were isolated by Histopaque-1077 density centrifugation. The separation and culture of naive CD4+ T cells were conducted according to our previous study^16^. The culture conditions used in this study were as follows: HSP condition – cell cultured with IL-15 and -7 (10 μg/ml, PeproTech); HSP+TCR condition – cells cultured with IL-15 and -7 and activated by anti-CD3 (5 μg/ml, BD Biosciences) and CD28 (1 μg/ml, BD Biosciences) on day 12; TCR condition – cells activated with anti-CD3/CD28 antibodies on day 0 and cultured with IL-2 (2 ng/ml, PeproTech); TCR+TCR condition – cells activated with anti-CD3/CD28 on day 0, cultured with IL-2 and second time activated with anti-CD3/CD28 at day 12.

### RNA-sequencing and bioinformatic analysis

Total RNA from cultured cells was isolated according to the manufacturer’s instructions using Qiagen RNeasy Micro kit (Qiagen) and submitted to the Genomics Unit of Centre for Genomic Regulation (CRG, PRBB, Spain) for sequencing. Quality and concentration of RNA were determined by an Agilent Bioanalyzer. Sequencing libraries were generated by a Ribo-Zero kit (Illumina). cDNA was synthesized and tagged by addition of barcoded Truseq adapters. Libraries were quantified using the KAPA Library Quantification Kit (KapaBiosystems) prior to amplification with Illumina’s cBot. Four libraries were pooled and sequenced (single strand, 50nts) on an Illumina HiSeq2000 sequencer to obtain 50-60 million reads per sample. RNA-seq reads were mapped against the Homo sapiens reference genome (GRCh37.p13) with the GEMtools RNA-seq pipeline (http://gemtools.github.io/docs/rna_pipeline.html). Genes were quantified with the same pipeline using the Gencode version 19 as an annotation. Normalization was performed with the edgeR TMM method^75^. The k-means clustering was performed by iDEP. 94 interface^17^ and the heatmap was generated by ggplot2 R package. Gene ontology (GO) enrichment analysis was performed with DAVID (http://david.ncifcrf.gov/). Differentially expression analysis was performed with the ‘robust’ version of the edgeR R package^76^. Genes with FDR<5% were considered differentially expressed. The list of human gene families was obtained from the HUGO gene nomenclature committee (https://www.genenames.org/). The list of HIV-interacting genes and restriction factors were obtained from the NIH HIV interaction database^18-20^ and a previous report^28^, respectively.

### RNA isolation and quantitative PCR

Total RNA was extracted from cells and treated with DNase I according to the manufacturer’s instructions using Qiagen RNeasy Mini Kit (Qiagen). 100 to 1000 ng RNA was reverse-transcribed into cDNA in a total volume of 20 μl using SuperScript IV Reverse Transcriptase (ThermoFisher). 2 μl of cDNA was used for quantitative PCR (qPCR) in a 10 μl reaction using SYBR master mix (ThermoFisher). Each reaction was performed in triplicates in 348-well plates in a QuantStudio 12K flex (ThermoFisher). Relative RNA levels were calculated after normalization to *18S rRNA* unless specified otherwise. Primers used this study are described in Table S7.

For *SLFN12* expression quantification in HIV-1 participants, total RNA samples from available PBMC dry-pellets were obtained using Qiagen RNeasy Mini Kit (Qiagen) following manufactureŕs recommendations. cDNA was obtained using SuperScript IV Reverse Transcriptase (ThermoFisher) and TaqMan gene expression assay (Applied Biosystems) was used for detection of *SLFN12* (Hs00430118_m1) and *TBP* (Hs99999910_m1). Gene amplification was performed on an Applied Biosystems 7500 Fast Real-Time PCR System thermocycler, and the relative expression was calculated as 2−ΔCT (where CT is the median threshold cycle from 3 replicates).

Quantification of integrated HIV-1-proviral DNA was performed from PBMC by droplet digital PCR (ddPCR) as described^77^. Briefly, two different primer/probe sets annealing to the 5′ long terminal repeat (LTR) and *gag* regions, respectively, were used to circumvent sequence mismatch in the patient proviruses, and the *RPP30* housekeeping gene was quantified in parallel to normalize sample input. Raw ddPCR data were analyzed using the QX100™ droplet reader and QuantaSoft v.1.6 software (Bio-Rad).

### Plasmids

SLFN11/12 sequences were acquired from the plasmids clone MGS: 59997 (*SLFN11*) and clone MGS: 45076 (*SLFN12*) provided by Dharmacon and cloned into mCherry expression vector, pmCherry-N’ (Clontech) between NheI and HindIII restriction sites. The plasmids generated were named pmCherry-SLFN11 and pmCherry-SLFN12, respectively. pNL-E, the HIV-1 proviral clone pNL4-3^78^ -derived *EGFP*-expressing plasmid, was generated previously^50^. The plasmid expressing HIV-1 wild-type *gag* sequence in pEF-BOS_bsr backbone was produced previously (pGag-wt)^79^. To generate HIV-1 codon-optimized *gag* expressing plasmid (pGag-opt), the codon-optimized *gag* sequence was obtained from the plasmid p96ZM651gag-opt (Y. Li, F. Gao, and B.H. Hahn through the National Institutes of Health (NIH) Reagent Program) and ligated between BamHI and NotI sites of pEF-BOS_bsr. Retrovirus vectors carrying shRNA sequences were constructed by inserting annealed oligonucleotides into a pSIN-siU6 vector (Takara) between BamHI and ClaI sites. As a negative control pSIN-siU6 vector expressing shRNA scramble sequences was used^80^. The constructs expressing mutant SLFNs (SLFN11 E209A, SLFN11 E214A, SLFN12 E200A and SLFN12 E205A) were generated using Q5 Site-Directed Mutagenesis Kit (NEB). The mutagenesis was performed according to the substitution protocol using primers designed with the NEBaseChanger (NEB). Codon-swapped *EGFP* expression vectors were produced previously^59^. The primers and shRNA sequences utilized here are listed in Table S7.

### Transfection and transduction

Transduction and transfection were done according to our previous study^81^. To prepare recombinant retroviruses carrying the shRNAs, HEK 293T cells were seeded in a 6-well plate at a density of 6 × 10^5^ cells/well and co-transfected with pSIN-siU6 – shSLFN11/ shSLFN12/ shSc (2 μg) along with plasmids pGP (1 μg, Takara) and pPE ampho (1 μg, Takara) using 12 μl of Lipofectamine 2000 (ThermoFisher) per well. At 48h after transfection, the supernatant was harvested and added on to ACH2 cells (1 ml of supernatant/10^5^ cells). Transduction was performed by spinoculation (1200 ×g, 25°C, 2h). Pelleted cells were resuspended in 500μl of RPMI with 10% FBS and placed into 24-well plates. Selection of transduced cells was done by adding G418 (1 mg/ml, InvivoGen) into the cell at 48h after transduction.

For the expression studies in HEK 293T cells, the cells were seeded in 24-well plate at a density of 2 × 10^5^, and after 24h co-transfected with construct pmCherry-SLFN11/ pmCherry-SLFN12 along with vector HIV-1-pNL-E, pGag-wt, pGag-opt, or codon-swapped EGFP using Lipofectamine 2000 (2.5 μl/well). At 3-4h post transfection, fresh medium was replaced, and the cells were kept at 37°C with 5% CO_2_ for 24 or 48 h.

### SAHA treatment of ACH cells

ACH2 cells with blocked expression of SLFN11 and SLFN12 (as described above) were seeded in a 24-well plate and treated with DMSO (0.01%, Sigma-Aldrich) or SAHA (Vorinostat, Sigma-Aldrich) at 0.5μM. The cells were lysed for RNA or protein isolation at 48h after treatment. Supernatants were harvested at 72h after treatment to test for virus titer by TZM-bl assay.

### TZM-bl assay

Virus titers in supernatants from transfected HEK 293T cells and HIV-1-reactivated ACH2 cells were determined by the TZM-bl assay as described previously^82^. Briefly, 11 serial dilutions of supernatants were prepared and added into fresh TZM-bl cells (10^4^ cells/well) in 96-well flat-bottom culture plates (Greiner Bio-One). After 72h incubation at 37°C, 5% CO_2_ the luciferase activity was measured on Centro LB 960 Microplate Luminometer (Berthold Technologies) using Britelite Plus^TM^ (PerkinElmer) according to the manufacturer’s protocol. Based on the luciferase levels, the TCID_50_ was calculated.

### Western Blot and antibodies

The cells were lysed with 1x passive lysis buffer (Promega), snap-frozen in liquid nitrogen and kept at -80°C overnight. The next day, cell debris was removed by centrifugation at 15,000 RPM for 5 min and supernatants were mixed with 2× Laemmli buffer, heated at 97°C for 5 min, transferred to the nitrocellulose membrane and immunoblot using specific antibodies was performed. Detection was done using secondary antibodies conjugated with horseradish peroxidase (HRP). Protein bands were developed on Medical X-Ray Blue Films (AGFA) using Pierce^TM^ ECL Plus Western Blotting Substrate or SuperSignal WestFemto Maximum Sensitivity Substrate (ThermoFisher) and quantified in Image-J software. The antibodies used in this study are listed in Table S8.

### Flow cytometry

CD4+ T cells after HSP or TCR culture were incubated with LIVE/DEAD® Fixable Aqua (ThermoFisher) and human Fc blocker (BD Bioscience). Then the cells were washed and stained with anti-CD4, CD8, CD27, and CD45RO- antibodies (Table S8) on ice for 20 mins. Transfected HEK 293T cells were suspended with PBS containing 1% FBS and 1μg/mL 4’,6- diamidino-2-phenylindole (DAPI) before data acquisition. Fluorescence was measured on a BD LSRFortessa (Becton, Dickinson) and analyzed with FlowJo v10 (Becton, Dickinson).

### Polysome profiling

HEK 293T were co-transfected with pmCherry/ pmCherry-SLFN11/ pmCherry-SLFN12 (12 μg) and HIV-1-pNL-E (12 μg) using Lipofectamine 2000 (48 μl) in a T-150 culture dish (ThermoFisher). In order to freeze elongation ribosomes, 48h after transfection cells were treated with 10 ml of DMEM containing cycloheximide (CHX, 100 μg/ml) during 2 min at 37°C and washed with 10 ml of PBS containing CHX (100 μg/ml) using a vacuum system. Cells were lysed with 700μl of lysis buffer (10 mM Tris-HCl (pH=7.4), 10 mM MgCl_2_, 100 mM NaCl, 1% Triton X100, 2 mM DTT, 100 ug/ml CHX), scraped and immediately frozen in liquid nitrogen and stored at -80°C. Cell lysates were thawed at 25°C and centrifuged at 12,000×g, 5min, 4°C and the supernatants were transferred to new tubes. After quantification, aliquots of 8 UA_260_ were made and stored at -80°C. Linear gradients of 10-50% sucrose were prepared in polysome buffer (20 mM Tris-HCl (pH=7.4), 10 mM MgCl_2_, 100 mM NH_4_Cl). The Gradient Master (Biocomp) was used to prepare the gradients in polyallomer tubes (Beckman Coulter). One aliquot of 8 UA_260_ was loaded on each gradient and centrifuged in Beckman SW41 rotor at 35 000 RPM, 3h, 4°C. Gradients were fractionated with fraction collector Model 2128 (BioRad). These fractions were used for phenol:chloroform RNA extraction and analysed by RT-qPCR.

### Structural comparison

Structure of human SLFN11 (predicted by alpha-fold, AF-Q7Z7L1-F1), SLFN12 (7LRE) and SLFN13 (predicted by alpha-fold, AF-Q68D06-F1) was obtained from Uniplot. A docking analysis was performed between SLFN12 and a human tRNA for Selenocystein (3HL2) according to Yang et al^49^.

### Calculating and visualizing gene-wise CAI and RSCU

For all hg38 RefSeq genes, we used the seqinr (v.3.3-6) and ggplot2 (3.1.0) R-packages and codon weights obtained from CAIcal (http://genomes.urv.es/CAIcal/CU_human_nature). We used Codon Usage Table database (https://hive.biochemistry.gwu.edu/cuts/about) for calculation of RSCU^55^.

### FISH-Flow

PBMC from ART-treated HIV-1-infected patients were obtained from the HIV Unit of the Hospital Universitari Vall d’Hebron (Barcelona, Spain) and CD4+ T cells were isolated by negative selection using magnetic beads (MagniSort Human CD4+ T Cell Enrichment; eBioscience). CD4+ T cells were then stimulated during 22h with latency-reversing agents (LRAs; 40 nM Romidepsin (Selleckchem) plus 100 nM Ingenol-3-angelate (Sigma-Aldrich)), or the negative control (media alone, R10; RPMI medium supplemented with 10% FBS). Prior to viral reactivation, cells were pre-incubated with the pan-caspase inhibitor Q-VD-Oph for 2h. In order to block new rounds of viral infections during viral reactivation, cells were treated with LRAs in the presence of Raltegravir (1µM), Darunavir (1µM) and Nevirapine (1µM). Cells were then subjected to the RNA FISH/flow protocol for the detection of HIV-1 transcripts and the viral protein Gag-p24 following the manufacturer’s instructions (Human PrimeFlow RNA Assay; eBioscience) with some modifications, as previously described^83^. In these experiments, to identify CD4+ T cells expressing SLFN12, HIV-1 RNA and the viral protein p24, the following antibodies were used: for cell surface staining, CD3 (PE-Cy7, BD Biosciences); and for SLFN12 detection, after fixation and permeabilization steps, cells were stained with primary rabbit anti-SLFN12 (Abcam) followed by incubation with a secondary donkey anti-rabbit IgG (AF488, Invitrogen) for 30 min at room temperature. The expression of HIV-1 RNA transcripts was analyzed with HIV-1 *gag-pol*-specific AF647-labelled probes, and the expression of the Gag- p24 viral protein was detected with a PE-anti-p24 antibody (clone KC57 RD1; Beckman Coulter). Cell viability was determined using an aqua viability dye for flow cytometry (LIVE/DEAD® Fixable Aqua Dead Cell Stain kit, Invitrogen).

### Statistical analysis

Student’s t-test, one-sample t-test, ratio paired t-test, Mann-Whitney test, or Spearman’s rank test (specified in each figure legend) were applied for the study. p-values <0.05 were considered statistically significant.

### Patient information

Classification of HIV-1 infected individuals was according to their HIV-1 RNA copies/mL (pVL) plus their history of ART in the past year. The criteria were as follows: (i) HIV-1 high viremic patients (HIV-High); >50.000pVL, (ii) HIV-1 low viremic patients (HIV-Low); <10.000pVL without ART, (iii) elite controllers (EC); <50pVL without ART, (iv) Viremic controllers (VC); <2000pVL without ART. The pVLs and CD4 counts of each patient group are summarized in Table S9.

## Acknowledgements

We thank Dr Song Gao for providing SLFN13-tRNA structure information, and Dr Maria-Eugenia Gas Lopez and Dr Ester Gea-Mallorquí for advise. This work was supported by following grants: MKI, JSPS Oversea Research Fellowship and Takeda Science Foundation; AEC, PT17/0009/0019 (ISCIII/MINECO and FEDER); MJB, RTI2018-101082-B-I00 and PID2021-123321OB-I00 [MINECO/FEDER]), and the Miguel Servet program by ISCIII (CP17/00179 and CPII22/00005); CB,MRR, CDC: European Union’s Horizon 2020 research and innovation program under grant agreement 681137-EAVI2020 and NIH grant P01- AI131568; JD, the Spanish Ministry of Science and Innovation (PID2019106959RB- I00/AEI/10.13039/501100011033; AM, the Spanish Ministry of Science and Innovation grant no. PID2019-106323RB-I00 AEI//10.13039/501100011033 and the institutional “María de Maeztu” Programme for Units of Excellence in R&D (CEX2018-000792-M.

## Author Contributions

MKI, JPM, YTY, JA, JD and AM designed the research study. MKI, KFS, JPM, YTY, FEA, KG, MG, JGE, CDC, and SR performed the experiments. JJ, ML, MD, and MY provided essential tools. AEC, RB, BO, MRR, and MJB analyzed the data. MKI, KFS, and AM wrote the manuscript. JA, JPM, CB, and JD critically discussed the manuscript. All authors have read and approved the final manuscript.

## Supplementary Methods

### Cytoplasmic and nuclear RNA isolation

HEK293T cells were transfected with pmCherry/ pmCherry-SLFN11/ pmCherry-SLFN12 along with HIV-pNL-E. 24h after transfection, the cells were lysed with lysis buffer (10 mM Tris-HCl (pH=7.5), 10 mM NaCl, 1.5 mM MgCl2, 10 mM Vanadyl complex (NEB), 1% NP- 40 (Sigma-Aldrich) and kept on ice for 5 min. After centrifugation at 3000 RPM for 5 min at 4°C, the supernatant (cytoplasmic fraction) was transferred to a new tube and the nuclear pellet was resuspended in a lysis buffer without Vanadyl complex and MgCl2. To extract RNA from each fraction, an equal volume Roti-Aqua-Phenol (Carl Roth, Germany) was added and centrifuged at 16000 xg for 5 min at room temperature. The aqueous phase was transferred into a new tube and subjected to chloroform extraction and ethanol precipitation. The ethanol precipitation was done by adding 0.3 volumes of 3M Sodium Acetate, 3 volumes of 100% EtOH (Merck) and 1 μl of glycogen (Roche). After 15 min incubation at -80°C, samples were centrifuged for 15 min at 16000 xg of 4°C. The supernatant was discarded and the pellet was resuspended in 1 ml of 70% EtOH, and proceeded to centrifugation for 5 min at 16000 xg of 4°C. The RNA pellet was dissolved in Diethyl Pyrocarbonate (DEPC)-treated water, and followed by TURBO DNA-free kit (Ambion) for DNA removal. Expression levels of HIV-gag RNA and HIV total RNA were analyzed by RT-qPCR.

### RNA gel electrophoresis

Transfected HEK 293T cells were collected and subjected to total RNA extraction through phenol-chloroform extraction as described above. 2µg of RNA were resolved in a 10% TBE-Urea polyacrylamide gel. Bands of tRNA and 5.8S rRNA were visualized by ethidium bromide staining and quantified by Image J.

**Figure S1.**
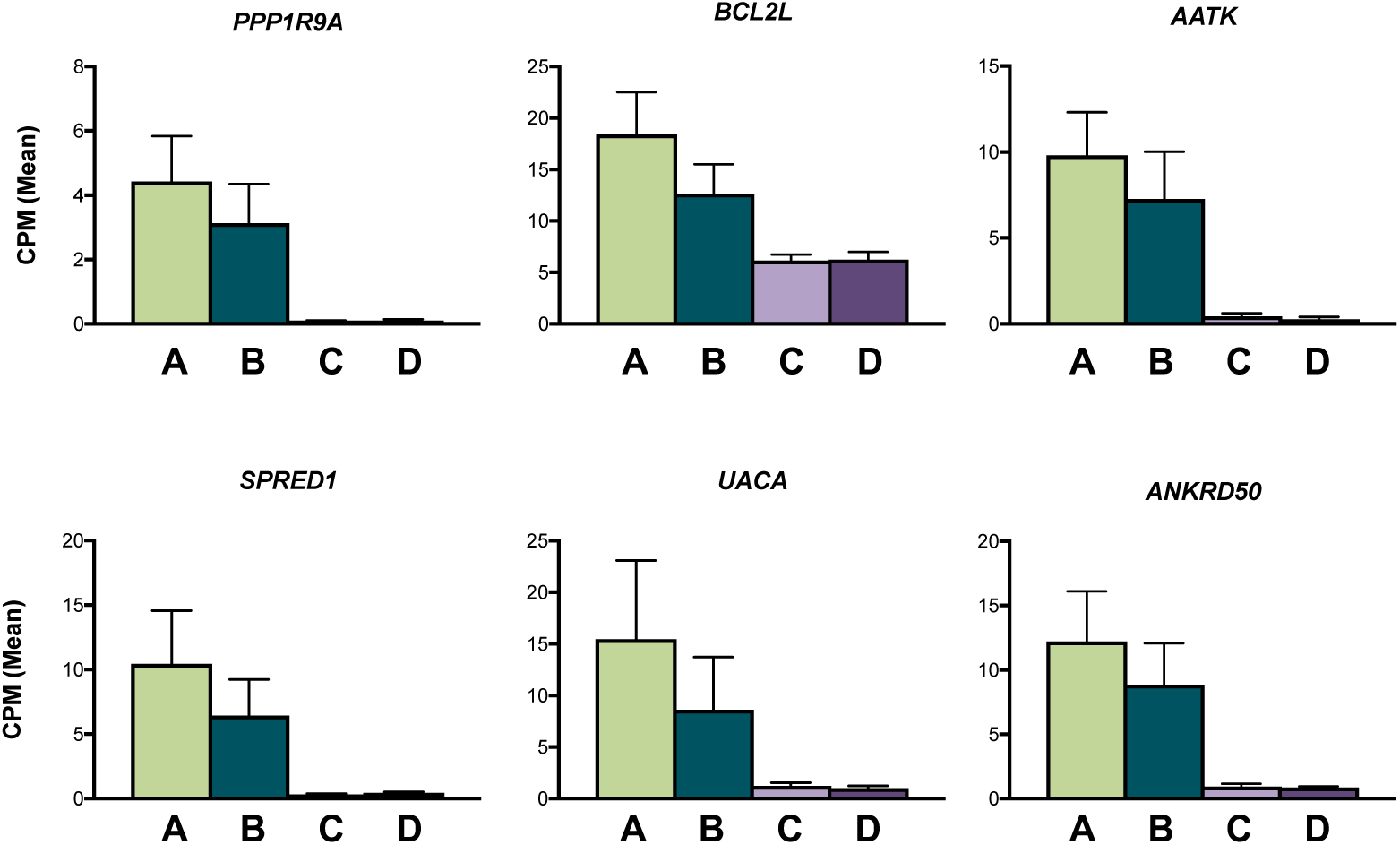
Expression patterns of the rest six genes in the final candidate list. Plots show mean CPM ± SEM (n=3).

**Figure S2.**
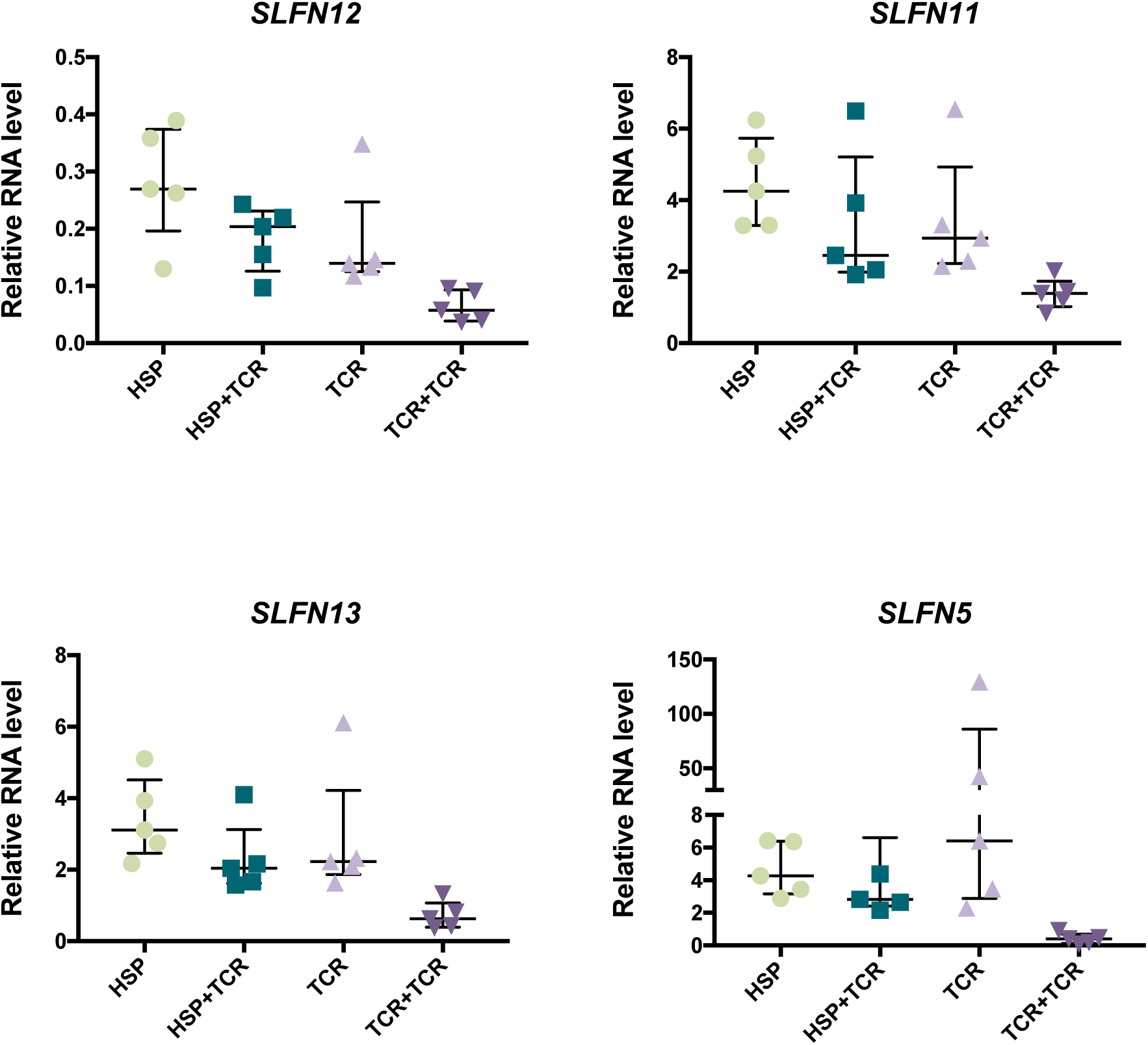
RT-qPCR data of five additional healthy donors cultured with HSP or TCR conditions. The bars show the median ± interquartile range (n=5). *SLFN14* was under the detectable limit in all the samples we tested.

**Figure S3.**
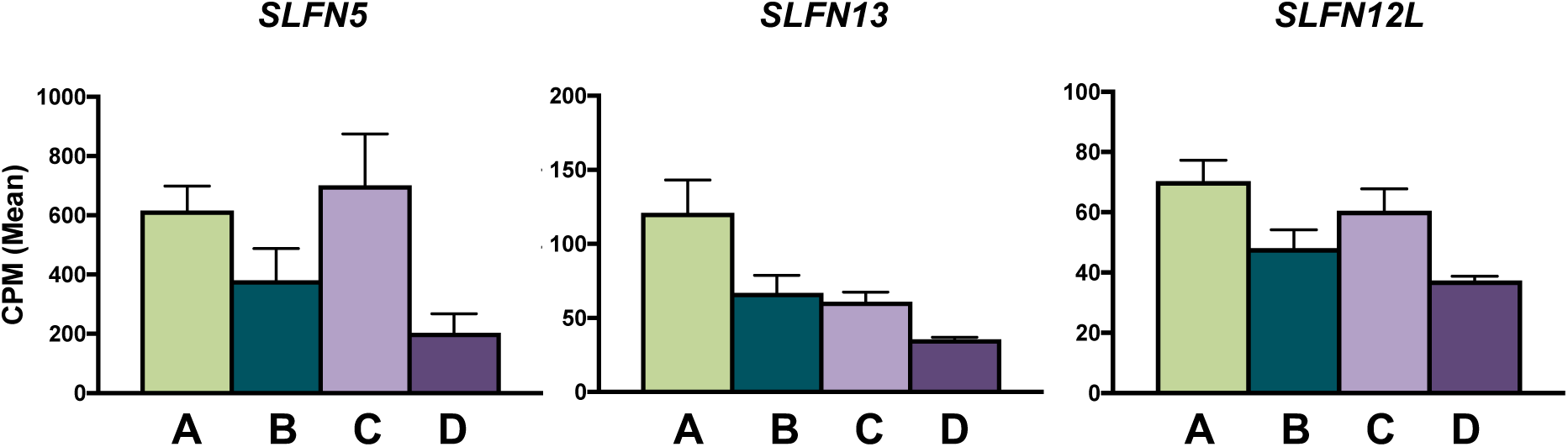
Expression patterns of *SLFN* family genes except *SLFN11 and SLFN12*. Given are the mean CPM ± SEM (n=3). *SLFN14* and *SLFNL1* showed no read count. *SLFN13* and *SLFN5* were classified into cluster II by the k-means clustering shown in Fig. 1c.

**Figure S4.**
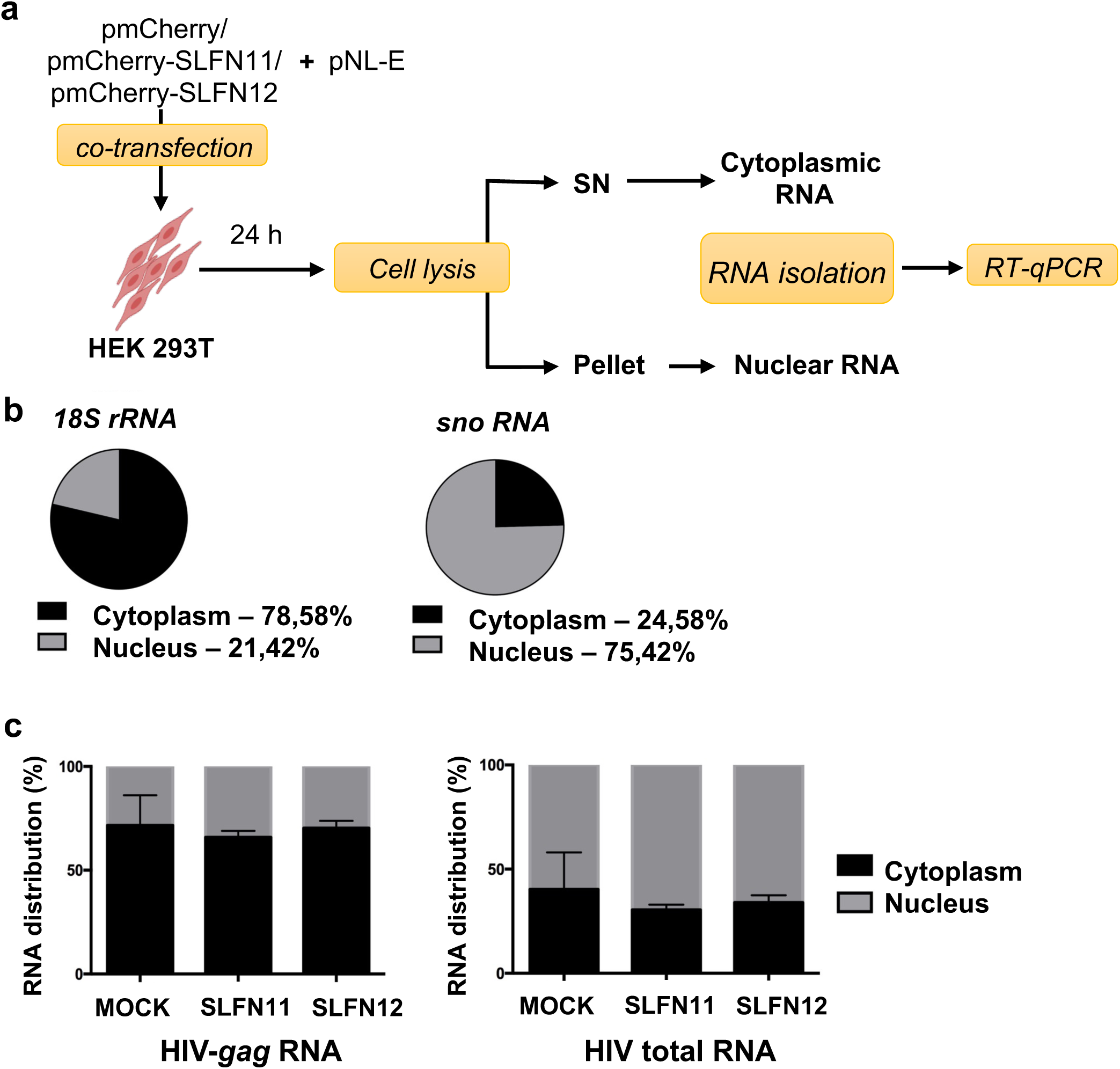
SLFN12 do not affect HIV RNA distribution. **a.** Scheme of this experiment. HEK 293T cells were co-transfected with pmCherry-SLFN11/12 or pmCherry empty vector along with HIV pNL-E vector. 24 hrs post-transfection, the cells were lysed in a hypotonic buffer to separate cytoplasmic fraction (supernatant) and nuclear fraction (pellet). RNAs were extracted from each fraction and analyzed by RT-qPCR. **b.** Fractionation efficiency. *18S rRNA* and *U3 small nucleolar RNA* (*snoRNA*) were quantified as controls for cytoplasmic and nuclear RNAs, respectively. Cytoplasmic and nuclear RNA distribution shown in percentage was calculated as an average of independent triplicates from mock-transfected HEK 293T cells. **c.** Distribution of HIV-*gag* RNA (Left) and HIV total RNA (Right) in the transfected HEK 293T cells. The total amount of HIV-*gag* or –total RNA in both fractions (cytoplasmic and nuclear) was set to 100%. Error bars represent the standard deviations of three independent samples.

**Figure S5.**
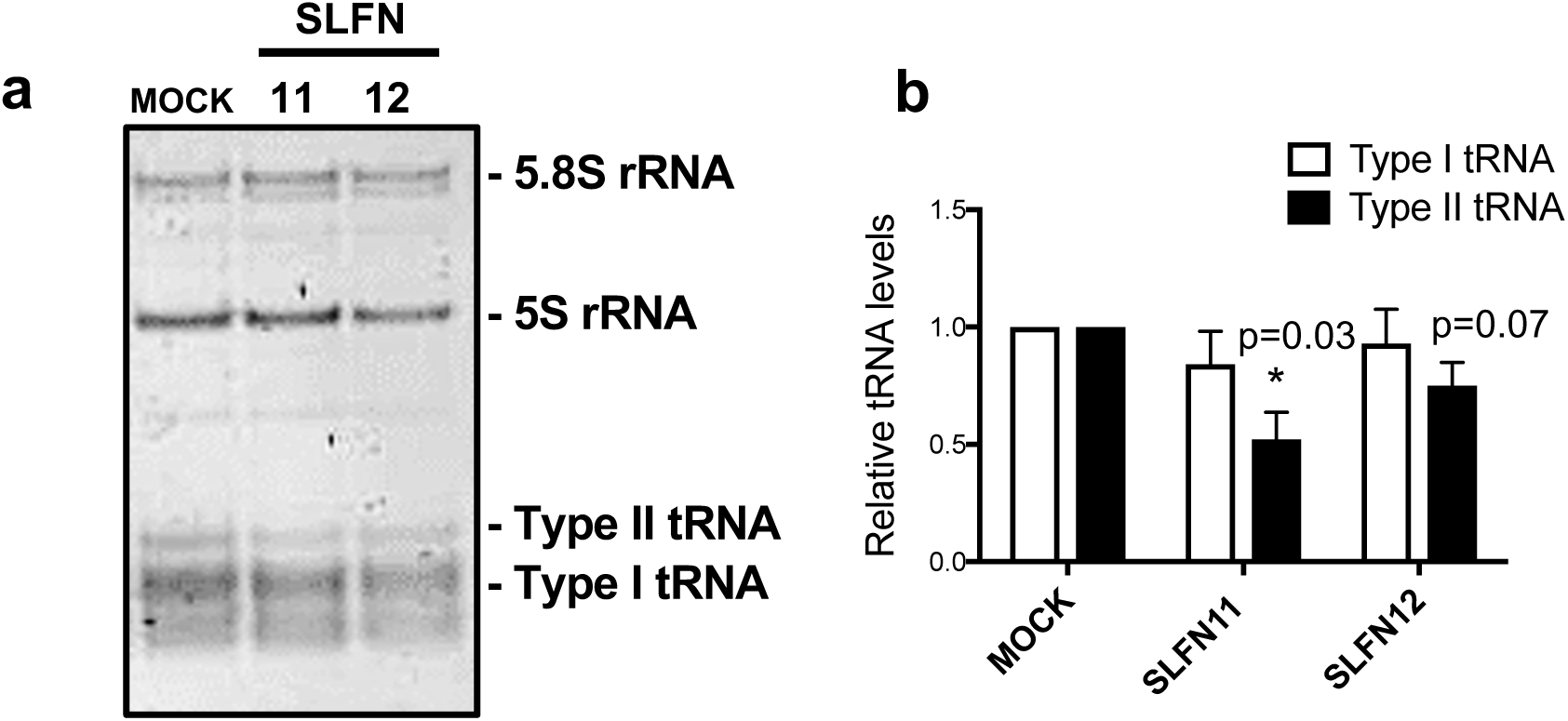
tRNA levels in cells after SLFN11 or SLFN12 expression. **a.** Representative result of RNA electrophoresis. Total RNA extracted from mock/SLFN11/SLFN12-expression vector-transfected cells were resolved on a 10% denaturing 10M urea polyacrylamide gel. **b.** Quantification of results from **a.** Plots show intensities of the type -I and -II tRNA measured by Image J and normalized by 5.8S rRNA intensity (n=3, mean ± SD). The levels in mock-transfected cells were set to 1. p-values were calculated by one-sample t-test.

## List of Supplementary Tables

Table S1. CPM values of each samples obtained from the RNA-seq

Table S2. Gene list shown up by k-means clustering

Table S3. List of known inhibitors of HIV-1 replication/function in the cluster I according to the NIH HIV interaction database

Table S4. List of DEGs and members of gene families that contain a known restriction factor (RF)

Table S5. List of the 58 candidate genes identified Fig. 1f

Table S6. Raw data of RSCU shown in Fig. 5e

Table S7. Oligos used in this study

Table S8. Antibodies used in this study

**Table S9.**
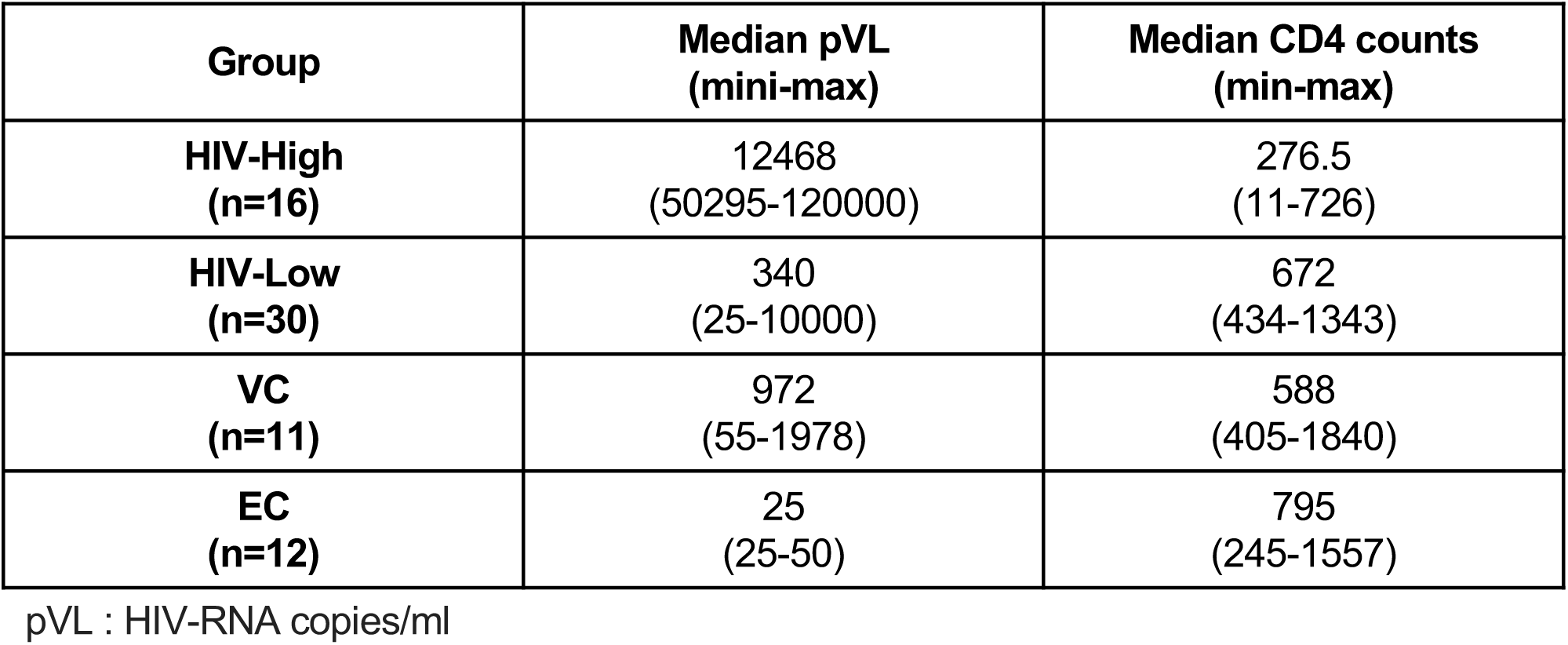
Clinical information of the patients used in Figs. 7a-c

